# Curli amyloid fibers in *Escherichia coli* biofilms: the influence of water availability on their structure and functional properties

**DOI:** 10.1101/2022.11.21.517345

**Authors:** Macarena Siri, Agustín Mangiarotti, Mónica Vázquez-Dávila, Cécile M. Bidan

**Affiliations:** Max Planck Institute of Colloids and Interfaces, Department of Biomaterials, Potsdam, Germany; Max Planck Institute of Colloids and Interfaces, Department of Sustainable and Bio-inspired Materials, Potsdam, Germany

**Keywords:** Amyloid fibers, Curli, Biofilm matrix, Bio-sourced materials, Spectroscopy

## Abstract

*E. coli* biofilms consist of bacteria embedded in a self-produced matrix mainly made of protein fibers and polysaccharides. The curli amyloid fibers found in the biofilm matrix are promising versatile building blocks to design sustainable bio-sourced materials. To exploit this potential, it is crucial to understand i) how environmental cues during biofilm growth influence the molecular structure of these amyloid fibers, and ii) how this translates at higher length scales. To explore these questions, we studied the effect of water availability during biofilm growth on the conformation and functions of curli. We used microscopy and spectroscopy to characterize the amyloid fibers purified from biofilms grown on nutritive substrates with different water contents, and micro-indentation to measure the rigidity of the respective biofilms. The purified curli amyloid fibers present differences in the yield, structure and functional properties upon biofilm growth conditions. Fiber packing and β-sheets content correlate with their hydrophobicity and chemical stability, and with the rigidity of the biofilms. Our study highlights how *E. coli* biofilm growth conditions impact curli structure and functions contributing to macroscopic materials properties. These fundamental findings infer an alternative strategy to tune curli structure, which will ultimately benefit to engineer hierarchical and functional curli-based materials.

**Graphical Abstract:** 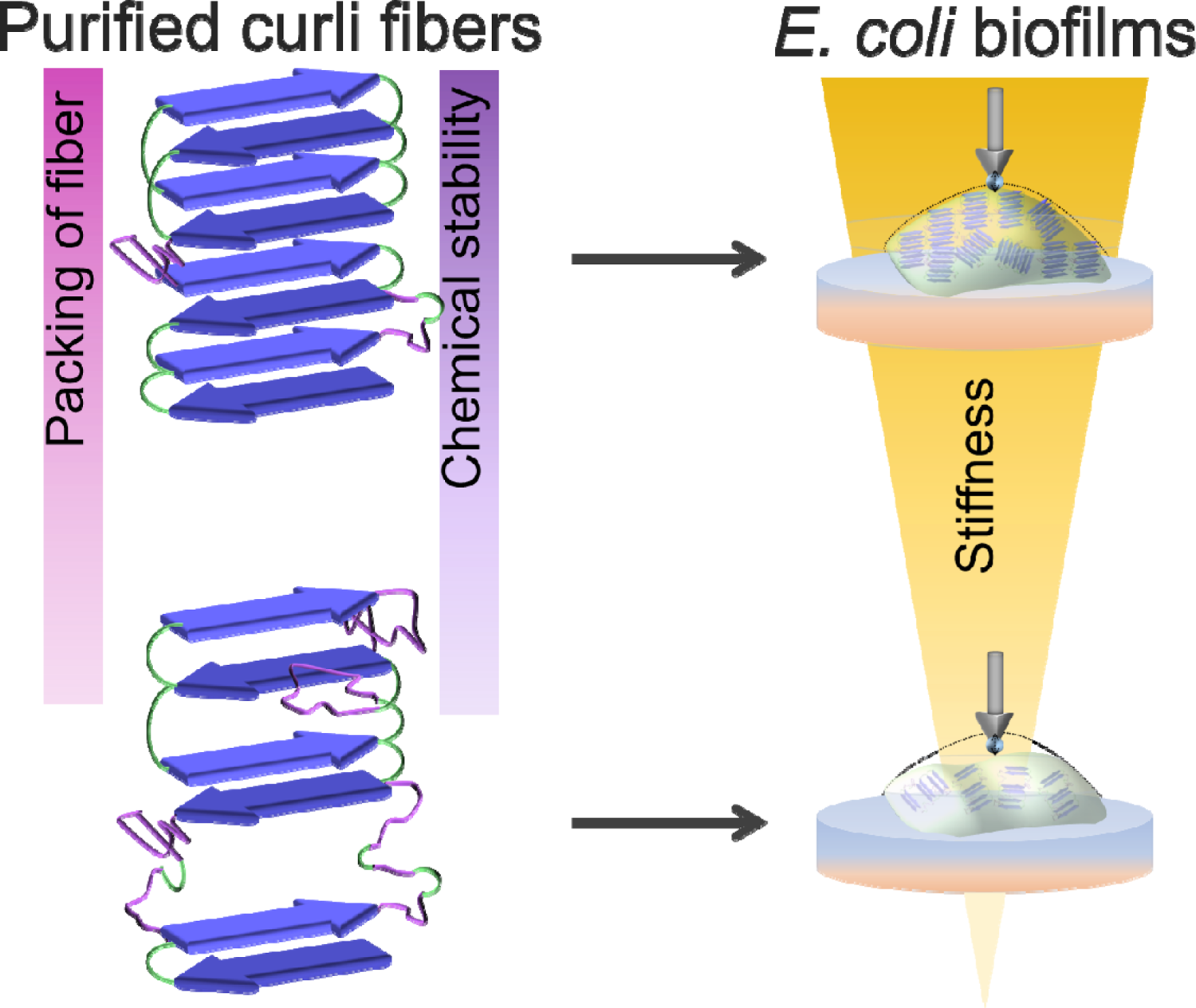

## Introduction

Biofilms are surface-associated groups of microbial cells that are embedded in a self-produced extracellular matrix. These living materials provide bacteria with resistance against environmental stresses such as exposure to antibiotics.^1–3^ In biofilms, the matrix consists of a network of biopolymers, mainly made of polysaccharides, proteins, and nucleic acids.

Amyloid fibers are often found in biofilm proteinous matrix. For example, *Bacillus subtilis* biofilms contain amyloid fibers assembled from the protein TasA, *Staphylococcus aureus* biofilms contain amyloid fibers made of phenol-soluble modulins (PSMs) and *Pseudomonas* biofilms contain fibers made of functional amyloids called Fap.^4^ In the last decade, the curli amyloid fibers made of CsgA proteins and found in *Escherichia coli* (*E. coli*) biofilms caught the attention of researchers aiming at engineering living and/or bio-sourced materials.^5^ Indeed, curli are particularly rich in β-sheets, which provide them with a notable rigidity and a remarkable stability that is further supported by the interactions of their side chains.^6^ These features have proved to yield attractive functional properties through their structural and protective roles in the bacterial biofilms,^7,8,9^ where they form tangled networks in a basket-like manner around the cells.^3^ Moreover, *E. coli* is a common biofilm-forming bacterium that can be found in the human digestive system and has been domesticated as *E. coli* K-12 for wide lab uses, so that it is now easily amenable to genetic modifications.^2,7^ As such, synthetic biologists have engineered *E. coli* to produce functionalized curli amyloid fibers,^10^ which further serve as building blocks to make materials ranging from living hydrogels to sustainable plastics and conductive composites.^10–13^

Because bacteria are living entities that adapt to their environment, their metabolism and matrix production activity are sensitive to their surroundings. Consequently, the shape, structure and materials properties of a biofilm depend not only on the organisms considered,^14,15^ but also on their growth conditions.^14,16^ For instance, the presence of salts, oxidative stress, surface charges and/or the water content of the substrate influence biofilm morphogenesis *via* different mechanisms.^3,7,16–20^ For example, the flux of water driven by osmotic gradients induces biofilm swelling and nutrient transport, which in turn governs bacteria proliferation and matrix production.^7,16,18^ Moreover, the confinement of bacteria growth by the substrate influences biofilm density, gives rise to complex biofilm morphologies and thereby modulates bacterial access to nutrients. These processes have been proposed to determine several properties of the biofilms,^7,19,20^ including their mechanical properties.^8,14,15,17^

In addition to the numerous studies focusing on the macroscopic scale, much *in vitro* work has been done to understand the assembly and aggregation kinetics of curli in aqueous solutions.^5,7,21^ For this, a collection of biophysical techniques, namely circular dichroism, infrared spectroscopy and fluorescence spectroscopy, have proved to be powerful tools to characterize amyloid fibers.^5,7,21,22^ Nevertheless, it is not still clear whether and how environmental cues experienced by the bacteria during biofilm growth influence the molecular structure of the matrix fibers. Yet, this piece of fundamental knowledge would be of great interest not only to further leverage the full potential of curli amyloid fibers as tunable building blocks to make bio-sourced materials, but also to develop strategies to prevent biofilm formation where they are detrimental, e.g. in medical and industrial contexts.^11,12,23^

The present work aims at bridging this gap by studying the effect of water availability during *E. coli* biofilm growth on the conformation and the functions of curli. For this, we cultured biofilm-forming *E. coli* bacteria of the strain W3110 on nutritive substrates with different water contents.^3^ After 5 days of growth, we purified the curli fibers from the biofilms obtained in each condition and performed a comparative study of the final fiber conformation using well-established microscopy and spectroscopy techniques. This approach enabled us to demonstrate that the overall mass, the structure, and the properties of the curli fibers assembled in the *E. coli* biofilms depend on the availability of water during growth. Indeed, curli amyloid fibers purified from biofilms grown on substrates with low water content presented higher hydrophobicity and chemical stability than those obtained from biofilms grown on substrates with high water content. Moreover, the changes in the structure of the matrix fibers reflected the changes in the biofilm mechanical properties. These results highlight the versatility of the curli fibers and their key role in the adaptation of biofilms properties to the physicochemical cues of their environment. As such, this work will greatly benefit to the research aiming at engineering biofilm properties and use their matrix components as key build blocks to make programmable living and/or bio-sourced materials.

## Results

To investigate the influence of water availability on the assembly of curli fibers in biofilms, *E. coli* from the strain W3110 were grown on salt-free Lysogenic Broth (LB) plates prepared with different agar contents, namely 0.5%, 1.0%, 1.8%, and 2.5 % agar.^17^ These small variations in the substrate composition lead to large variations in their properties, which in turn served as different environments for biofilm growth (**Figure S1**). In contrast to the *E. coli* strain AR3110 used in previous works and that produces a matrix made of both curli and phosphoethanolamine cellulose fibers,^17^ the strain W3110 only produces curli fibers. A brief analysis of the macroscopic features (i.e. size, mass, water content and water uptake upon rehydration) was first conducted on the biofilms obtained in the different growth conditions. A purification protocol (see Experimental Section) was then applied to the different biofilm samples to harvest the curli fibers from the matrix. Finally, detailed structural and physico-chemical characterizations of the curli fibers were performed using microscopy and spectroscopy techniques and micro-indentation was used to assess the rigidity of the corresponding biofilms.

### Substrate water content influences biofilm size, mass and composition

After 5 days of growth, the biofilms were observed under transmission light with a stereomicroscope (**Figure 1a**). The biofilms cultured on 0.5 % salt-free LB-agar (i.e. on wet substrates) are significantly larger compared to biofilms grown in the other conditions (i.e. on dryer substrates), while biofilms grown on 1.0 % salt-free LB-agar did not differ in size compared to the standard condition (1.8 % salt-free LB-agar). Weighing the biofilms after harvesting them from the surface showed that the bacteria cultured on substrates with lower agar contents formed biofilms of significantly higher wet mass than on substrates with higher agar concentrations (**Figure 1b**). We further estimated biofilm water contents, which revealed no significant differences between biofilms grown in the different conditions (**Figure 1c**). They only spanned from 74.11 % (biofilms grown on 2.5 % salt-free LB-agar) to 77.18 % (biofilms grown on 0.5 % salt-free LB-agar) with a relatively large variability. Upon overnight rehydration of dried samples, the biofilm material grown on 0.5 % salt-free LB-agar showed the lowest water-uptake (**Figure 1d**).

**Figure 1.**
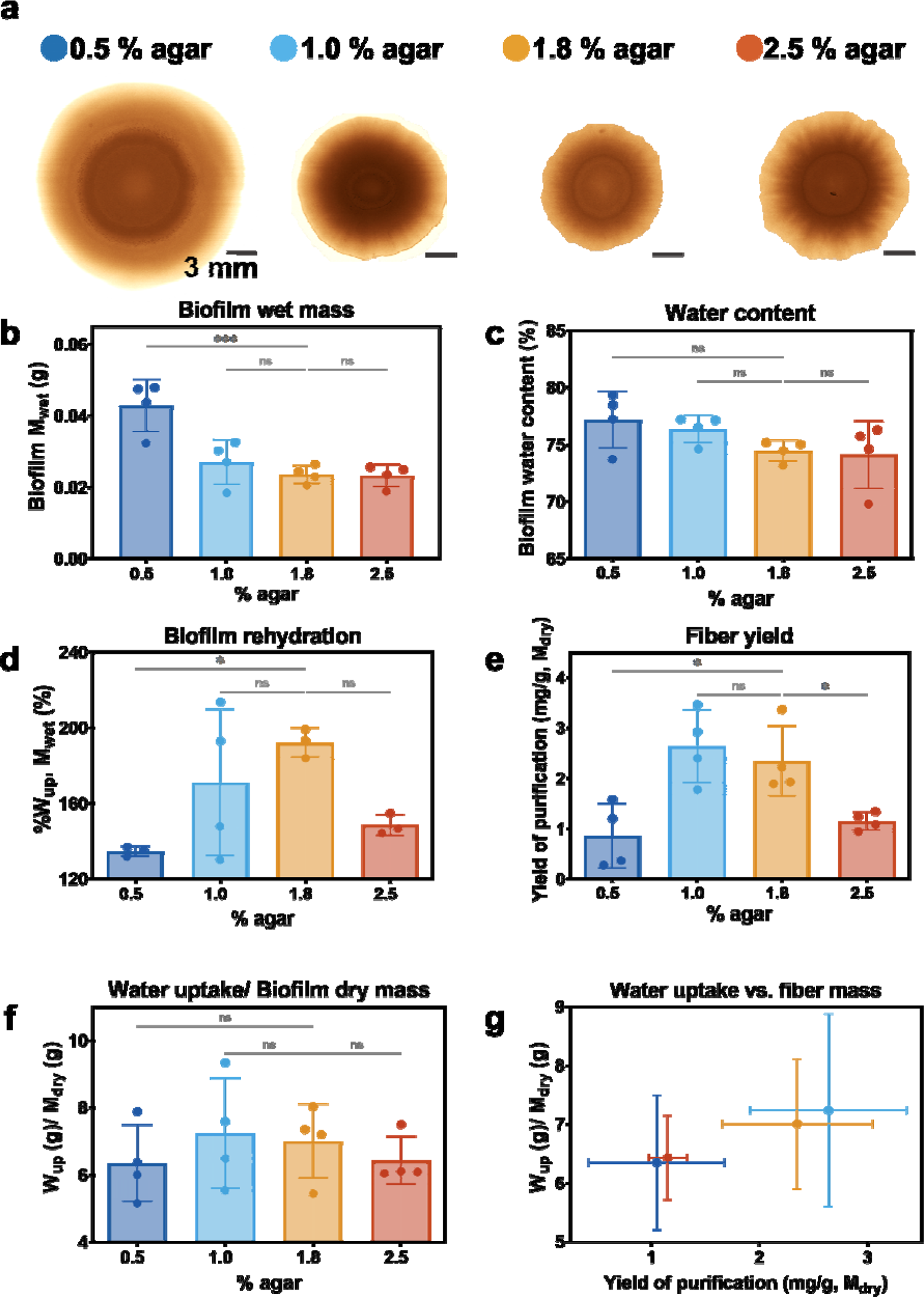
General characteristics of *E. coli* W3110 biofilms grown on salt-free LB-agar substrates with different water contents. (a) Representative phenotype of five-day old biofilms grown on salt-free LB-agar substrates with different water contents. (b) Biofilm wet mass after scraping them from the surface. (c) Water content of biofilms calculated as W = (m_wet_−m_dry_)/m_wet_ × 100% w/w. (d) Water uptake per biofilm after overnight rehydration in excess of water calculated as %W_up,w_ = (m_rewet_ – m_dry_)/m_wet_ × 100% w/w. (e) Purification yield of the curli fibers extraction process in milligram of CsgA per gram of biofilm dry mass. The CsgA content in the purified curli fibers was estimated by absorbance after denaturation using 8 M urea. (f) Water uptake per gram of dry biofilm calculated as W_up,d_ = (m_rewet_ – m_dry_)/m_dry_. (g) Water uptake as a function of fiber mass (i.e. yield) per gram of dry biofilm in each condition. All data presented here come from N=4 independent biofilm cultures for each condition tested, and the statistical analysis was done with One-way ANOVA (p<0.001, *** | p<0.01, ** | p<0.05, * | ns = non-significant).

A standard purification process^21^ was then applied to the biofilms grown on the substrates with different agar contents (i.e. different water contents) in order to isolate the curli fibers secreted by the bacteria to form the matrix (see Experimental section). The purification yield, defined as the mass of curli fibers per gram of dry biofilm, was then estimated from the concentration of CsgA monomers measured by absorbance after treatment of the fibers with urea (**Figure 1e**). Interestingly, the highest purification yield was obtained from biofilms grown on substrates containing 1.0% and 1.8 % agar, i.e. the standard condition used to grow biofilms in previous studies involving *E. coli.*^3,18^ Both the highest and the lowest agar concentrations tested led to biofilms containing less curli fibers. Indeed, biofilms grown on 0.5 % and 2.5 % salt-free LB-agar yielded 2 times less curli fibers than those grown in standard conditions (1.8 % salt-free LB-agar). Note that this tendency follows the one obtained on the water uptake per gram of dry biofilm after rehydration (**Figure 1f**), where the dry mass is mostly made of dry curli and bacteria. Despite the presence of other molecules that can take up water in the biofilms (e.g. colanic acid), the similarity in the trends observed for the amount of curli produced by the biofilm and for the water uptake capacity of the matrix suggests an interesting role of curli fibers in biofilm swelling and water storage (**Figure 1e** and **f**).

To estimate how the water content of the substrate influences biofilm composition, we considered the whole mass harvested from these *E. coli* W3110 biofilms to be the sum of the masses of i) the water, ii) the curli fibers and iii) the bacteria, the remaining nutrients and other matrix biomolecules in minor proportions. The sum of the two latter is the dry mass of the biofilm (**Table 1**). Although all the growth conditions rendered biofilms with similar distributions of wet and dry mass (Figure 1b and **Figure S2**), there are some differences in the content of their dry masses (**Table 1**). Biofilms grown on dryer substrates (2.5 % salt-free LB-agar) present slightly lower content of dry mass, while those grown on wet substrates (0.5 % salt-free LB-agar) present higher content of dry mass. Both conditions, 0.5 % and 2.5 % salt-free LB-agar, yield biofilms with lower curli content in their dry mass composition compared to the other growth conditions.

**Table 1.** Composition of *E. coli* W3110 biofilms grown on salt-free LB-agar substrates with different water contents. The total mass corresponds to the sum of the water content and the dry mass of the biofilms. ^a^The mass of “bacteria + rest” was estimated by subtracting the curli mass, which is given by the quantification of CsgA monomers after purification from the dry mass. The percentages of curli and bacteria + rest are given with respect to the dry mass. N=4 independent biofilm cultures for each condition tested (27 biofilms per experiment).

| Growth condition | Units | Total mass | Water | Dry mass | Curli<sup>a</sup> | Bacteria + rest<sup>a</sup> |
| --- | --- | --- | --- | --- | --- | --- |
| 0.5 % agar | mg | 43.73 ± 7.31 | 33.04 ± 5.44 | 9.83 ± 2.27 | 0.008 ± 0.006 | 9.82 ± 2.27 |
|  | % | 100 | 77.18 | 22.82 | 0.085 | 99.91 |
| 1.0 % agar | mg | 27.04 ± 6.21 | 20.70 ± 4.96 | 6.34 ± 1.27 | 0.016 ± 0.003 | 6.33 ± 1.27 |
|  | % | 100 | 76.37 | 23.63 | 0.264 | 99.74 |
| 1.8 % agar | mg | 23.58 ± 2.53 | 17.57 ± 2.00 | 6.01 ± 0.57 | 0.014 ± 0.004 | 6.00 ± 0.57 |
|  | % | 100 | 74.46 | 25.54 | 0.235 | 99.77 |
| 2.5 % agar | mg | 23.25 ± 3.11 | 17.29 ± 2.85 | 5.96 ± 0.40 | 0.007 ± 0.001 | 5.95 ± 0.40 |
|  | % | 100 | 74.11 | 25.89 | 0.115 | 99.88 |

### Curli fibers formed in biofilms grown on wet substrates contain less β-sheet structures

To make sure that the purification yielded the expected product, the purified materials were examined by transmission electron microscopy (TEM), by fluorescence confocal microscopy after Thioflavin S (ThioS) staining and by circular dichroism (CD) and attenuated total reflectance Fourier transform infrared (ATR-FTIR) spectroscopy (**Figure S3** and Figure 2). The TEM images first confirmed that the purified samples show an amyloid-like morphology with needle-like structures (**Figure S3**). ThioS is a fluorescence probe that reports the presence of β-sheet pleated structures.^24^ As the samples stained positive for ThioS, the amyloid nature of the fibers was confirmed (Figure 2a).^24^ The differences in the ThioS intensity of the fibers suggested that biofilm growth conditions render fibers with different conformations (Figure 2a). In all the samples, agglomerated structures were detected and could be interpreted as clumps of tightly associated fibrils and protofibrils. These clumps remained after extended sonication prior to imaging, indicating strong adhesive forces linking the aggregates together.^5^ Despite not presenting significant differences, the intensity of the ThioS signal (relative to background) in the samples obtained from biofilms grown on 0.5 % salt-free LB-agar was slightly lower than in samples obtained from the other conditions (Figure 2a). To further assess the differences in fibers structure, we studied conformational characteristics by fluorescence (Figure 2b), CD (Figure 2c), and ATR-FTIR spectroscopy (Figure 2d-e).

**Figure 2.**
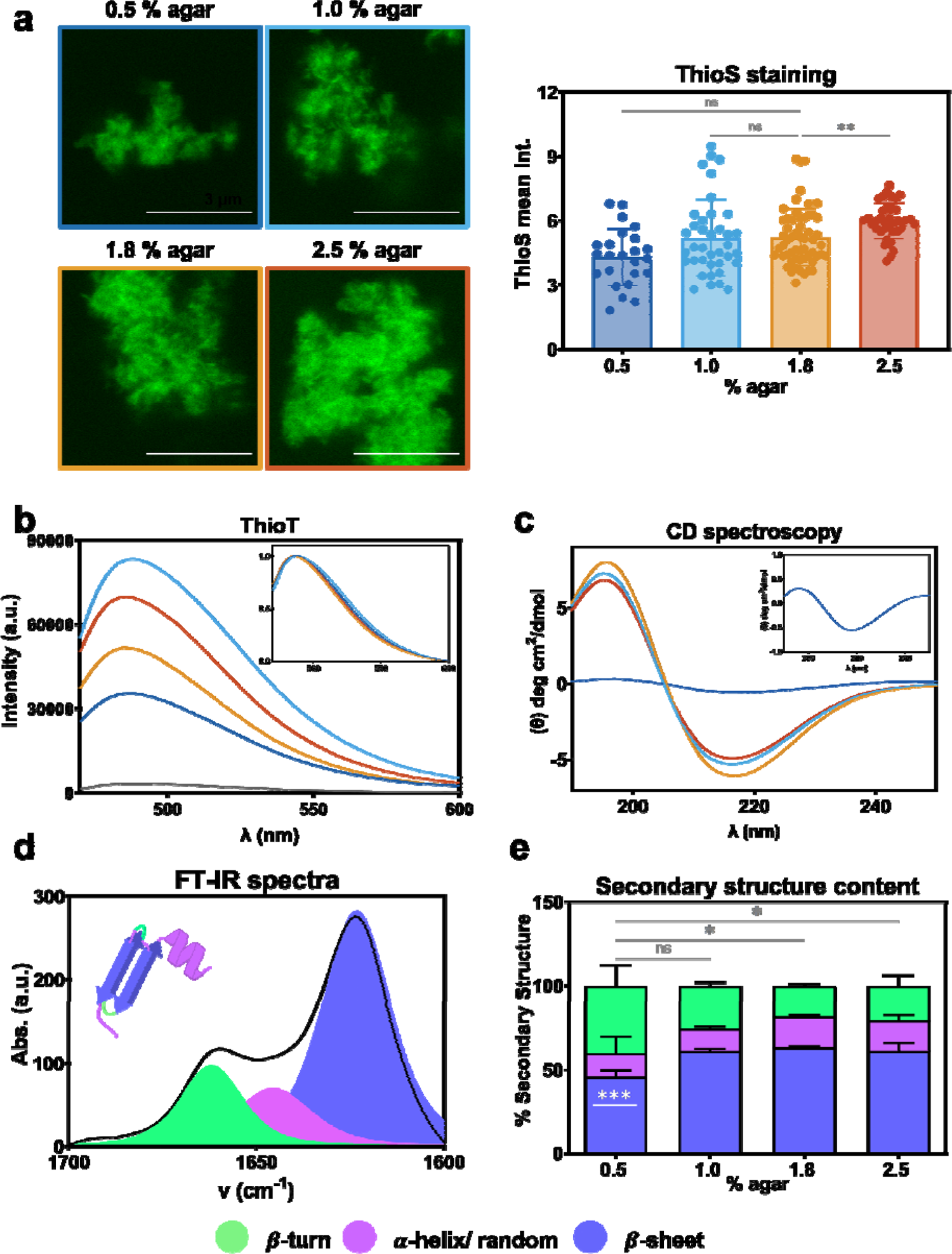
Characterization of curli fibers purified from *E. coli* biofilms. (a) Confocal images of purified curli fibers stained with ThioS (left) and corresponding mean fluorescence intensities (right). For statistical analysis Kruskal-Wallis test was used (p<0.01, ** | ns = non-significant) with a Dunn’s post-test for multiple comparisons (alpha=0.05) comparing all samples against the condition 1.8 % agar. N=5 to 10 independent fiber solutions obtained for each condition. Scale bar= 3µm. Fiber solutions had a concentration of 5 µM protein monomeric units (see experimental section for more details). (b) ThioT fluorescence emission spectra of the fibers at equivalent mass concentrations. N=3-4 independent fiber solutions obtained for each condition. The curves plotted are a representation of the average of the measurements taken. Fiber solutions had a concentration of 5 µM protein monomeric units (see experimental section for more details). (c) Circular Dichroism (CD) spectra of the fiber solutions obtained in different conditions. (Inset) zoom of the CD spectra corresponding to fibers obtained from biofilms grown on 0.5 % salt-free LB-agar. The curves plotted are an average of 3 measurements of the different fiber solutions. Fiber solutions had a concentration of 5 µM protein monomeric units (see experimental section for more details). (d) Representative ATR-FTIR absorbance spectrum obtained on curli fibers purified from *E. coli* biofilms grown on 1.8 % salt-free LB-agar, focused on the amide I’ region. The spectra were fitted (color curves) using the 3 spectral components identified by second derivative on the Fourier self-deconvoluted spectra (see Experimental section and Figure S4 for more details). (e) Distribution of the 3 types of secondary structure in the curli fibers obtained in the different biofilm growth conditions. For statistical analysis One-way ANOVA test was used (p<0.05, *) with a Dunnett’s post-test for multiple comparisons (alpha=0.05) comparing all samples against the condition 0.5 % agar. β-turns proportions (green) are significantly different when compared to fibers obtained from biofilms grown on a 0.5 % salt-free LB-agar, β-sheet proportions (dark purple) in fibers obtained in dry conditions (1.0, 1.8 and 2.5 % agar) are also significantly different to those obtained in wet conditions (0.5% agar) (Asterix in white, p<0.001, ***). N=4 independent fiber solutions obtained for each condition.

In order to allow for reliable comparison in the following experiments, the fiber solutions were all normalized to 5µM of monomeric CsgA units. ThioT was added to the different solutions to study fiber conformations. Indeed, it has been demonstrated that ThioT fluorescence intensity relates to the number of exposed β-sheets and/or the spacing between them,^25^ therefore qualitatively reporting fibrils packing.^26,27^ It was also reported that changes in ThioT emission could also reflect differences in the electrostatic interaction between the dye and the fiber.^28^ Nonetheless, non-significant differences were observed among the fibers after measuring their ζ-potential in the buffer used for the ThioT binding experiment (50 mM glycine buffer pH 8) (Figure S4). Note that in water, the fibers have negative ζ-potential values, and turned positive in buffer, as expected when increasing pH. This result suggests that differences in the ThioT spectra emission cannot be attributed to electrostatic interactions. Here, all fiber samples showed a 10 to 30 fold increase of the ThioT intensity compared to the reference without fibers (Figure 2b). Fibers assembled in the biofilms grown on 0.5 % salt-free LB-agar present the lowest ThioT intensity, which indicates that the corresponding amyloid-like β-sheets differ in nature, extent and/or packing of their β-strands.^27^

CD spectroscopy showed the signature of β-sheet signaling for all fiber solutions, with a maximum at ∼195 nm and a minimum at ∼216 nm.^5^ The differences observed between the different spectra indicate changes in the secondary structure content of the purified curli fibers (Figure 2c). Such changes are more pronounced for the biofilms grown in wet conditions (0.5 % salt-free LB-agar) (Figure 2c**, inset**). ATR-FTIR spectroscopy further confirmed the presence of amyloid fibers and gave additional insights into their secondary structure. Indeed, this technique is widely used to study protein structure and is especially suited for amyloids.^27,29,30^ The analysis was focused on the amide I′ band (from 1700 to 1600 cm^−1^) that mostly represents the amide C=O stretching and is especially sensitive to protein secondary structure.^29–31^ All the samples showed spectra similar to Figure 2d. The predominant band (Figure 2d, purple) was centered around 1620 cm^-1^ and assigned to β-sheet structures.^31^ Indeed, a narrow and intense absorption band between 1615 and 1630 cm^-1^ is a hallmark for amyloid fibers as it indicates a large, planar and extremely well-ordered cross-β spine.^29,30,32^ The second band located around 1660 cm^-1^ is usually assigned to β-turns structures (Figure 2d, green).^5,6,31^ Finally, using band fitting to decompose the spectra into its overlapping components systematically revealed a minor contribution around 1650 cm^-1^, which is assigned to random and α-helix structures (Figure 2d, pink).^29^ In the case of curli fibers purified from *E. coli* biofilms, the contribution of each of these two components overlapped and could not be distinguished.

Despite the similarities among the samples, the detailed analysis of the secondary structure contents in the amide I’ band showed significant differences in the packing of the curli fibers harvested from biofilms grown on the different agar substrates (Figure 2e **and Figure S4**). For example, band decomposition reveals that fibers formed in biofilms grown on substrates with high water content (i.e. 0.5% agar) contain below 50 % of β-sheet while the fibers grown in the other conditions reach 60 %. However, the curli fibers from biofilms grown on 0.5 % salt-free LB-agar contain almost twice the amount of β-turns structures compared to the other fibers.

### The hydrophobicity of the curli fibers depends on the water content of the biofilm substrate

To further detail the properties of the curli fibers obtained from biofilms grown in different conditions, we used the solvatochromic dye 1-anilinonaphthalene-8-sulfonic acid (1,8-ANS), which probes the local nano-environment around its moiety. 1,8-ANS emits a weak fluorescence signal that both increases and shifts its maximum upon binding to hydrophobic pockets.^33^ It is also a suitable dye to distinguish fiber polymorphism.^34,35^ In the presence of the fibers tested, 1,8-ANS showed a hyperchromic shift compared to the probe in buffer, and with a stronger shift for fibers obtained from biofilms grown in dry conditions (Figure 3a and **3b**). Phasor plot analysis helps to visualize the variations in the hydrophobicity of the fibers by means of the shift of the center of mass of each spectrum (Figure 3b and **S5**). In brief, phasor analysis consists in a Fourier transform of the spectral information (or lifetime, see Experimental section) into a two-dimensional space where the axes represent the real and imaginary components.^36,37^ In the case of spectral phasors, the angle carries the information of the spectral center of mass and the radial direction is related with the spectral width, providing a fingerprint of the system (See Experimental section and **Figure S5** for more details).^36,37^ As such, a visual inspection of the phasor plot allows to interpret the changes taking place between the systems, since points presenting different coordinates indicate that the dye is sensing a different nano-environment. In Figure 3b, the hydrophobicity increases clockwise, with 1,8-ANS in buffer being the least hydrophobic sample, as expected. Fibers grown in wet conditions have the lowest hydrophobicity, while those grown in standard conditions have the highest hydrophobicity as sensed by 1,8-ANS (Figure 3b). By calculating the ratio between the intensity of 1,8-ANS fluorescence emission in the fiber solutions and in buffer (Figure 3c), we observed that fibers obtained from biofilms grown on 1.0 and 1.8 % salt-free LB-agar present the highest increase by two-fold of 1,8 – ANS intensity (Figure 3c). Despite having non-significant differences, the trend for ThioT ratio variation for fibers in the different conditions matches the trend observed for 1,8 – ANS intensity variation (Figure 3d, derived from Figure 2c). The higher the packing of the fiber (ThioT ratio), the higher the hydrophobicity (1,8-ANS ratio) (Figure 3e). ThioT and 1,8-ANS have different targets within the fibers: ThioT is incorporated to the β-sheets,^25^ whereas 1,8-ANS binds to the exposed hydrophobic cavities of the fibers.^38,39^ These results suggest that the β-sheet content might be responsible for the hydrophobicity in the fibers.^5^

**Figure 3.**
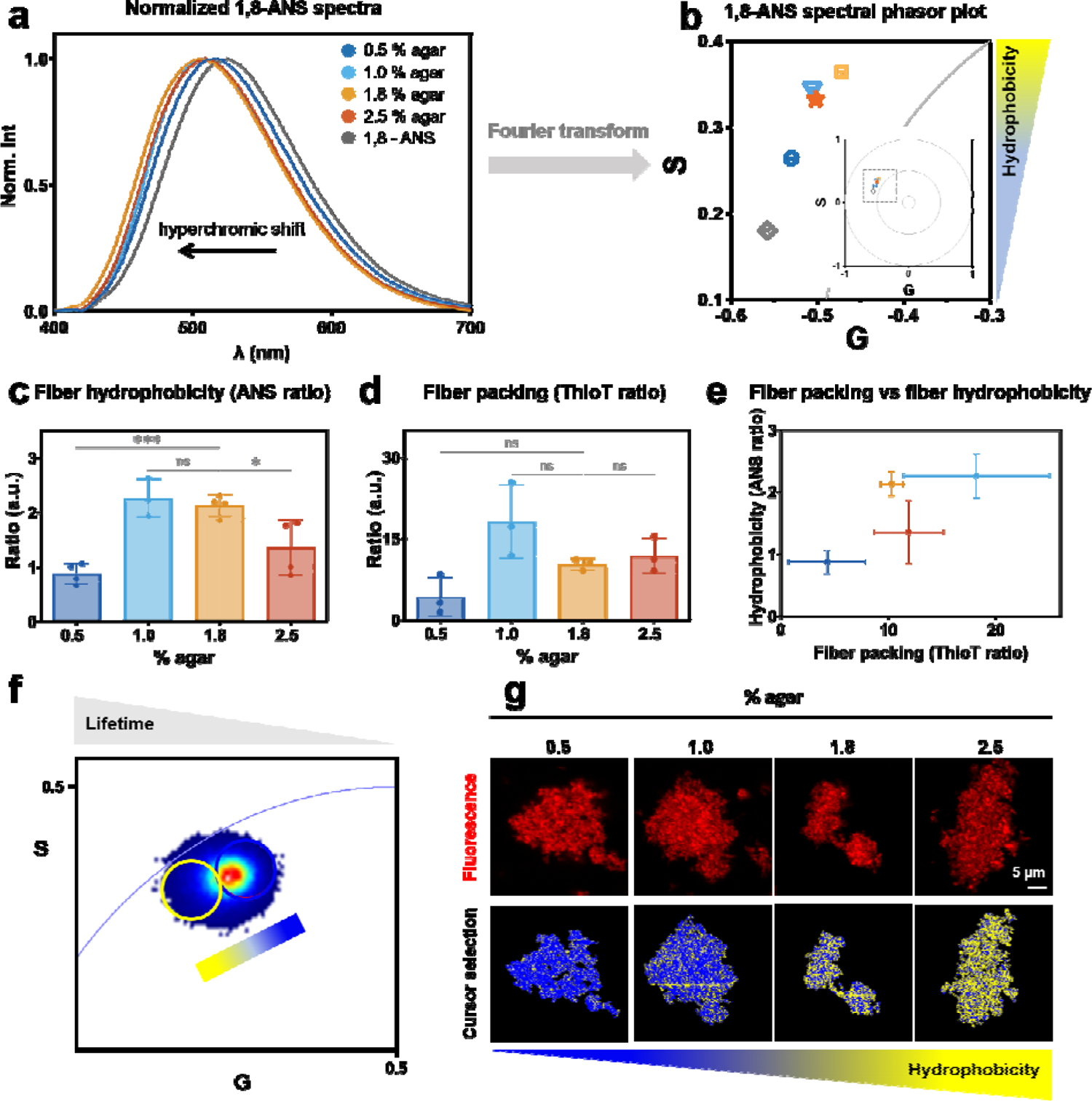
Hydrophobicity and packing of curli fibers purified from *E. coli* biofilms grown on salt-free LB-agar substrates of different water contents. (a) Normalized 1,8-ANS emission spectra in the presence of the fibers at equivalent mass concentration. Emission of 1,8-ANS in buffer is also included (grey). N=4 to 5 independent fiber solutions obtained for each condition. (b) Spectral phasor plot of the purified samples bound to the probe 1,8-ANS. S and G correspond to the real and imaginary parts of the Fourier transform of the spectra (see methods). Here hydrophobicity (hyperchromic shift) increases clockwise. (Inset) full plot where the dashed square indicates the magnified region. (c-d) Values describing the increase in intensity of the probes when bound to the purified fibers. Quantification of the increase was estimated by division of the area under the curve of each spectra of the probe (ANS or ThioT) with each fiber by the area under the curve of the emission spectra of the probe alone. For statistical analysis One – way ANOVA test was used (p<0.001, *** | p<0.05, * | ns = non-significant). N=4 to 5 independent fiber solutions obtained for each condition. (e) Correlation between the fiber hydrophobicity (ANS ratio) and fiber packing (ThioT ratio) using data from panel (c-d). (f) Nile Red FLIM phasor plot for the different purified fibers. Each pixel of the final image contains lifetime information. The data obtained from fibers formed in the different conditions are superimposed. The colors of the pixel clouds indicate the pixel density, increasing in a rainbow scale from blue to red. (see Figures S6 and S7 for more details). (g) Confocal microscopy images of purified fibers stained with NR (top) and phasor map (bottom) in which each pixel is colored according to the color of the corresponding cursor in the lifetime phasor plot in (f). The choice of the size and the position of the cursor is arbitrary and it is used to highlight average properties of the lifetime distributions (see Experimental section and S5). Increasing the agar content (i.e. decreasing water availability) during biofilm growth moves the pixel clouds towards longer lifetimes, i.e. increasing the hydrophobicity of the Nile Red binding site of the curli fibers. All fiber solutions had a concentration of 5 µM protein monomeric units (see experimental section for more details).

Nile red (NR) is another solvatochromic dye that has a high sensitivity for the tertiary structure of different amyloid fibers.^40^ NR has the advantage to be fluorescent at a visible wavelength (λ_em_=585 nm), making it convenient for fluorescence microscopy. Here we use a combined FLIM-phasor approach to assess the changes in NR lifetime within the fibers obtained in different biofilm growth conditions (see experimental section and S6).^41–43^ We show the representative confocal microscopy images of fibers stained with NR and the corresponding phasor plot for the lifetime of the fibers obtained from biofilms grown on the different substrates (Figure 3f-g). An increase in the lifetime of NR indicates a less polar, i.e. more hydrophobic environment at the binding site of the probe.^44,45^ By using colored cursors (blue and yellow circles), pixels of a given lifetime can be selected and the corresponding pixels are mapped back onto the image using the same color code (Figure 3f). In this case, the blue cursor corresponds to shorter lifetimes (more polar environments) and the yellow cursor to longer lifetimes (more hydrophobic environments). An increase in the agar concentration (i.e. decrease in water availability during biofilm growth) correlates with a higher lifetime of NR in the fibers (i.e. higher hydrophobicity, more yellow in Figure 3g). As such, changes in the NR lifetime, i.e. environment hydrophobicity, between the fibers obtained in different growth conditions can be easily identified without fitting procedures. A statistical quantification of these changes can be found in the supporting information (**Figure S7**).

### The tryptophan population of all fibers is accessible to the solvent

Considering the potential of curli as building blocks for bio-sourced materials, we aimed at getting more information on the supramolecular organization of the fibers. For this, we investigated their intrinsic fluorescence, which can be exploited to study changes in solvent accessibility of the side chains of aromatic amino acids.^46^ Curli is made of CsgA monomers, each one containing a single tryptophan (Trp), four tyrosine, and three phenylalanine residues.^32^ When curli is excited at 280 nm, the intensity of the intrinsic fluorescence emission is mainly due to Trp in CsgA.^46,47^ When we excite the purified fibers at 280 nm, a single maximum emission can be observed around 350 nm for all the spectra (Figure 4a). This position suggests that the environment of Trp is relatively polar and structureless. The data acquired is consistent with position of the Trp in the CsgA predicted by AlphaFold (Figure 4b).^47,48^

**Figure 4.**
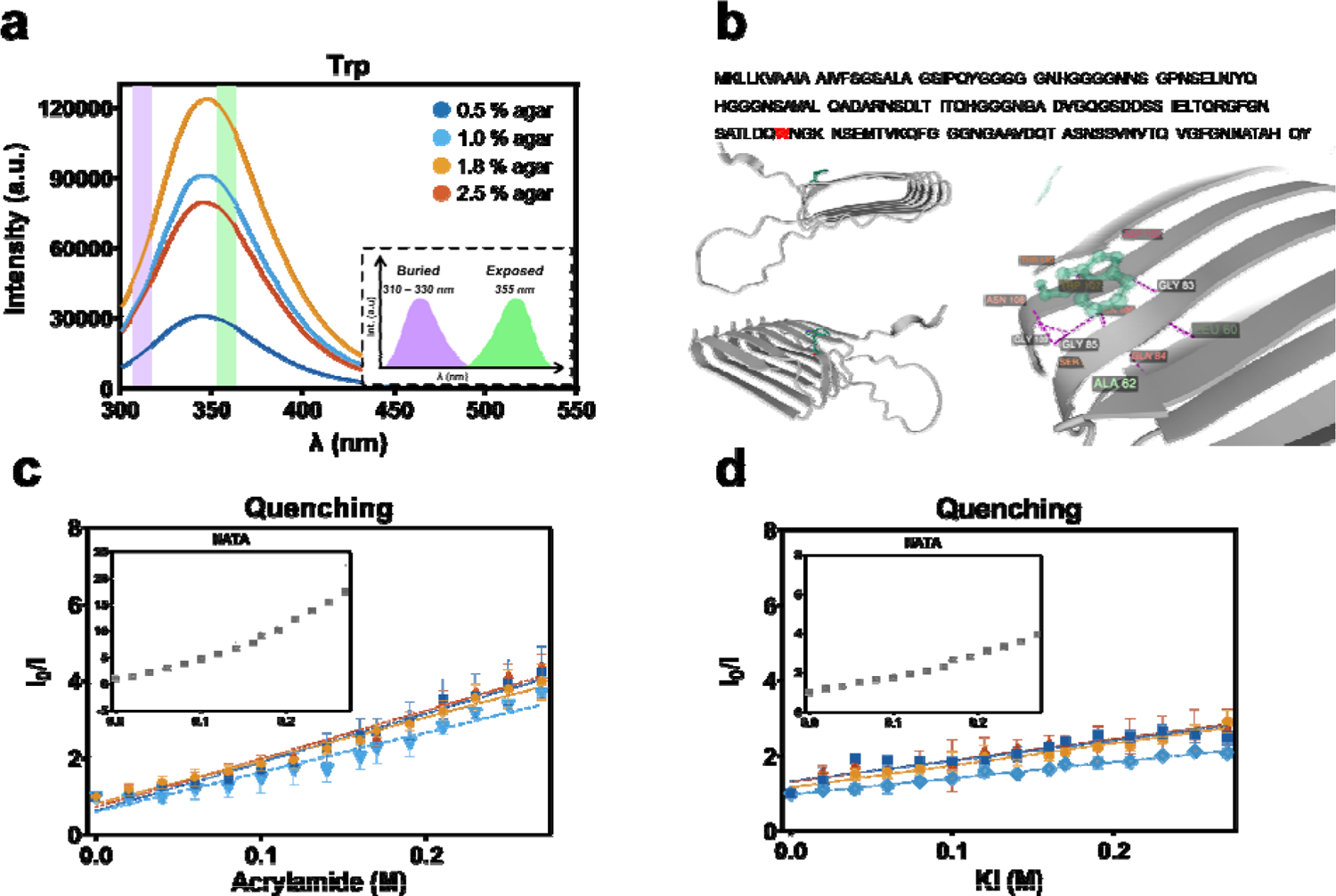
Characterization of the intrinsic fluorescence of the purified curli fibers. (a) Intrinsic Trp emission fluorescence spectra of the fibers at equivalent mass concentrations (λ_exc_ = 280 nm). A sketch is included as an inset to describe the expected position of the spectrum peak according to the Trp exposure.^50^ The shadowed areas in the main plot indicate these exposure extremes. (b) Position of the Trp residue in the CsgA protein as predicted by AlphaFold.^48^ The residues interacting with Trp^107^ are also depicted. The CsgA aminoacid sequence is shown with the Trp highlighted in violet. The sequence was obtained from Uniprot (Locus: A0A7H9LSK1). Stern-Volmer plots of (c) acrylamide or (d) iodide quenching of Trp residues in the purified curli fibers. I_0_ corresponds to the emission signal of ThioT incubated with fibers at the beginning of the experiment (urea concentration = 0M). (Inset) Effect of each quencher on NATA (grey). N=4 independent fiber solutions obtained for each condition. All fiber solutions had a concentration of 5 µM protein monomeric units (see experimental section for more details).

The steady-state emission spectrum of fibers assembled in biofilms grown in standard growth conditions (1.8 % salt-free LB-agar) have the highest emission intensities, whereas the fibers assembled in biofilms grown on substrates with high water content (0.5 % salt-free LB-agar) present the lowest Trp emission intensities. As explained above, the differences observed in the intensity of the purified fibers reflect changes in solvent accessibility of the side chains of aromatic amino acids. Hence, we explored the exposure of the Trp residues in each fiber using neutral and ionic dynamic fluorescence quenchers (Figure 4c-d), namely acrylamide (polar and uncharged) and iodine (highly hydrated, large and negatively charged).^49^ N-acetyl-L-tryptophanamide (NATA), a soluble version of tryptophan, was used as reference for the maximum quenching possible (Figure 4c-d **insets**).

The Stern–Volmer plots showed no upward curvature, meaning that the quenching effect in all cases was predominately dynamic quenching, i.e. that fluorescence deactivation occurs because of the collision of the quencher with the fluorophore and not because of a binding interaction between both molecules.^49,51^ Moreover, the addition of the quencher (acrylamide or iodide), even at high concentrations did not induce a shift in the spectra of the Trp in the purified fibers (**Figure S8**). This suggests that there is no alteration of the hydrophobic character of the fibers throughout the experiment.

Quenching experiments indicated no buried Trp residue within the fibers, suggesting that the Trp population of all the fibers studied is exposed to the solvent (Figure 4c-d).^52^ The plots of the ratio between the fluorescence intensities in the absence and the presence of quenchers (I/I_0_) as a function of the quencher concentration yielded straight lines under all external conditions within the concentration range 0–0.4 M. There are no significant differences for the value of the quenching slopes (K_SV_ values) of the purified fibers: all values are between 10 – 12 M^-1^ (**Table S2**). Of note, the K_SV_ values for acrylamide are much lower than those obtained with NATA denoting that the emitters are not exposed to the solvent in the fibers. This implies rapid diffusion of the quencher to the Trp.^49,52^

### Fiber chemical stability and biofilm rigidity follow fiber molecular structure

To study the implications of the structural differences described above, we tested the thermal and chemical stability of the purified fibers against exposure to a temperature ramp and different urea concentrations, but also the mechanical properties of the biofilms (Figure 5 and **Figure S9**).

**Figure 5.**
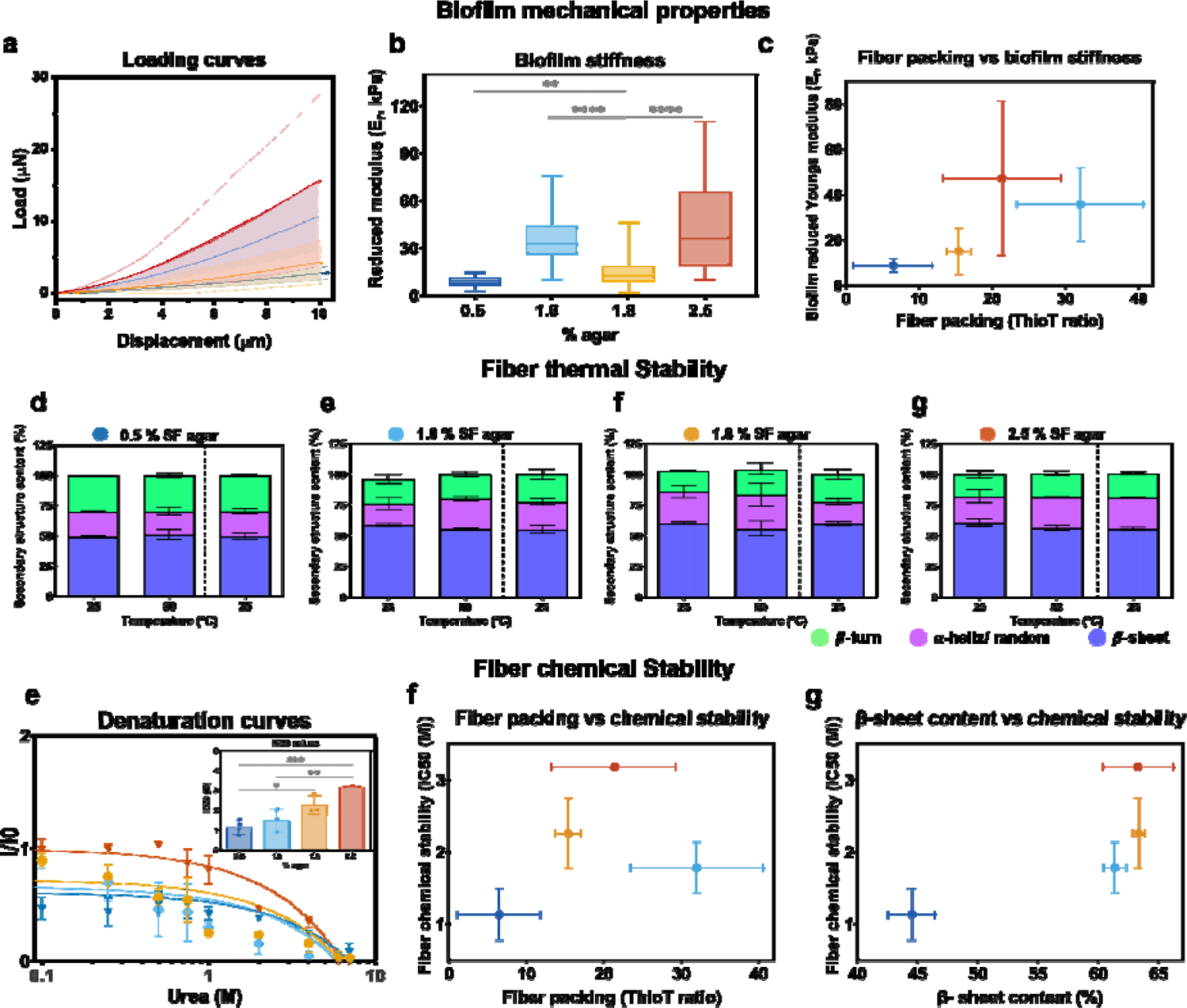
Functional properties of purified curli fibers and mechanics of the corresponding biofilms. (a-d) Thermal stability of fibers purified from biofilms grown on substrates of different water contents. Secondary structure contents by analysis of FTIR spectra of curli fibers treated with increasing temperatures (25 – 80 °C). The data after the dashed line represent the secondary structure contents of fibers after a sudden decrease of temperature (from 80 – 25 °C). N=3 fiber solutions per condition. (e) Chemical stability of the purified fibers upon denaturation with increasing urea concentrations (0.1 – 8 M). The presence of the fibers was observed by ThioT fluorescence emission intensity. I_0_ corresponds to the ThioT emission when bound to fibers without urea in the solution. (Inset) IC50 corresponds to the urea concentration at which the ThioT intensity is 50 % of the initial one. For statistical analysis One-way ANOVA test was used (p<0.001, *** | p<0.01, ** | p<0.05, * | ns = non-significant) with a Tukey’s post-test for multiple comparisons (alpha=0.05) comparing all samples against the 1.8 % salt-free agar concentration. N=4 fiber solutions per condition. (f-g) Fiber chemical stability as a function of (f) fiber packing or (g) fiber β-sheet content. (h) Loading curves of the micro-indentations performed on *E. coli* biofilms grown in each condition. (i) Biofilm reduced Young’s moduli obtained by fitting an Hertzian contact model the loading curves. n=5 to 10 curves were analyzed on N=3 to 4 biofilms per condition. For statistical analysis a Mann-Whitney U test was used (p<0.0001, **** | p<0.01, **). (j) Biofilm reduced Young’s moduli as a function of curli fiber packing for the respective biofilm growth conditions.

Thermal denaturation was performed by heating the fibers from 25 to 80°C and cooling them down back to 25°C (see Experimental section and Figure S9 for details). At each temperature step, the structure of the fibers was monitored using ATR-FTIR spectroscopy. All the samples of fibers showed a remarkable stability, with no significant change of structure as monitored by band analysis (Figure 5a-d).

The chemical stability of the fibers was tested against increasing urea concentrations from 0.1 to 8M (Figure 5e). The presence of amyloid fibers was monitored by the fluorescence intensity emitted by ThioT. The stability of the fibers significantly decreased as the water content of the corresponding biofilm the substrate increased (Figure 5e). We observed no linear correlation between the fiber chemical stability and its packing arrangement (Figure 5f). Nevertheless, we observed that the higher the β-sheet content in the fiber, the higher their chemical stability (Figure 5g).

Biofilm stiffness was assessed by micro-indentation experiments in the central area of the biofilms grown on different substrates with different water contents (**Figure S10**).^17^ For each biofilm, loading curves were obtained upon approximatively 10 nm indentation (Figure 5h). Although there is no clear trend, we observed that biofilms grown in wet conditions (0.5 % salt-free LB-agar) presented the lowest reduced Young moduli. Their stiffness even revealed to be significantly lower than the one of biofilms grown in other conditions. In contrast, biofilms grown on 1.0 % and 2.5 % salt-free LB-agar were significantly stiffer than those grown on 1.8 % salt-free LB-agar (Figure 5i). These results on the mechanical properties of the biofilm could be understood as macroscopic consequences of the changes of the curli structure in the matrix. We thus plotted biofilm reduced Young moduli as a function of the structure of the curli fibers formed in each corresponding condition (Figure 5j). We found that the higher the fiber packing (ThioT ratio), the stiffer the biofilm.

## Discussion

Amyloid fibers extracted from the microbial biofilm matrix have recently become interesting building blocks for engineering living and/or bio-based materials.^11–13,23^ Previous work have showed the influence of bacteria growth conditions on the materials properties of biofilms, and the determining role of external conditions such as molecular crowding, temperature and pH on amyloid fibrillation and fiber conformation.^5,14,27,53^ Here, we focus on how an environmental cue experienced by *E. coli* bacteria during biofilm growth – namely water availability – influences the molecular structure and properties of curli. For this, we cultured biofilm-forming *E. coli* bacteria of the strain W3110 on nutritive substrates of different water contents.^3^ The results demonstrate that the physical-chemical properties of curli depend on the water availability in the biofilm growth environment. Indeed, CD, ATR-FTIR and fluorescence spectroscopy revealed differences in the yield of curli fibers purified from the biofilms, in the packing of these fibers, their hydrophobicity and their chemical stability. Moreover, micro-indentation experiments revealed a relation between these matrix fiber properties and the rigidity of the resulting biofilms.

Water, especially bound water (or hydration water), plays an important role during the different stages of protein aggregation in fiber formation, and therefore in the final conformation the fiber adopts.^54^ Both experiments and molecular dynamics simulations suggest that water favors the interactions between hydrophobic regions of the monomers or between polyglutamine sequences, and lead to structure variations when trapped inside proteins.^55^ In a crowded environment, protein hydration shells tend to change and amyloid proteins are more prone to aggregation.^27^ As shown by the binding differences to ThioS and ThioT, the packing and the β-sheet content of the curli fibers depend on the water content of the biofilm substrate (Figure 2b-e).^24^ These structural differences in the fibers are further supported by the differences observed in tryptophan (Trp) emission signal, which reflect differences in Trp interactions with surrounding groups such as threonine (Thr), aspartic acid (Asp), glutamine (Gln), and asparagine (Asn) (Figure 4b).^27,47^ Indeed, these polar and charged residues have the ability to quench Trp fluorescence. Hence, the lower intensity in the spectra of fibers obtained from biofilms grown in wet conditions (0.5 % agar) can be further explained by changes in the structure of these fibers.

The fibers obtained from biofilms grown on substrates with high water content (0.5 % agar) also showed lower hydrophobicity, as sensed by the 1,8-ANS and the NR probes (Figure 3).^56^ Interestingly, the phasor plot obtained from 1,8-ANS spectra revealed that the fibers purified from biofilms grown on 1.8 % salt-free LB-agar present more hydrophobic binding sites (Figure 3a-b). Moreover, the similar trends of the ThioT and the 1,8-ANS intensities as a function of growth condition, suggest that the β-sheet content can be correlated to the hydrophobicity in the fibers (Figure 3c-e).^5^ NR fluorescence studies confirmed that fibers obtained in wet conditions are less hydrophobic than fibers obtained in dry conditions (Figure 3f-g). The difference between the results obtained from the 1,8-ANS and the NR experiments on the 1.8 % salt-free LB-agar are more likely due to the different binding sites and affinities of the probes.^39,57,58^ Nonetheless, both results indicate clear differences between fibers fibrillated in dry and wet conditions.

Amyloids are highly ordered and arranged into intermolecular β-sheets and cross β-structures, which contain many strong hydrogen bonds that enhance the overall stability of the fibers.^5,59^ Models of curli fibers also suggest that the hydrogen bonds between Gln and Asn residues from different monomers form a network that contributes to the stability of these fibers.^5,32,60^ Considering the importance of the structure-function relationship in proteins and the materials they form, the implications of the structural differences observed in the purified curli fibers were assessed with stability assays. While all fibers showed high stability against thermal denaturation,^9^ their stability upon urea exposure revealed significant differences (Figure 5h). The thermal stability can be attributed to the high content of β-sheets in fibers obtained from all conditions (Figure 2e). However, the relatively light packing and lower β-sheets content measured in curli fibers obtained at high water concentrations could explain their lower chemical stability compared to those obtained on dryer substrates (Figure 5h-i). Indeed, less densely packed fiber structures not only result in weaker interactions between the distant Gln and Asn residues, but they also provide better access for urea to denature the fibers by interactions with their hydrophobic and amide groups (Figure 6).^61^ Note that the fibers extracted from biofilms grown on 0.5% salt-free LB-agar not only contain less than 50% of β-sheet but almost twice as much β-turns compared to the other conditions (Figure 2e), which indicates that β-turn structure is less chemically stable than β-sheet (**Figure S11**).

**Figure 6.**
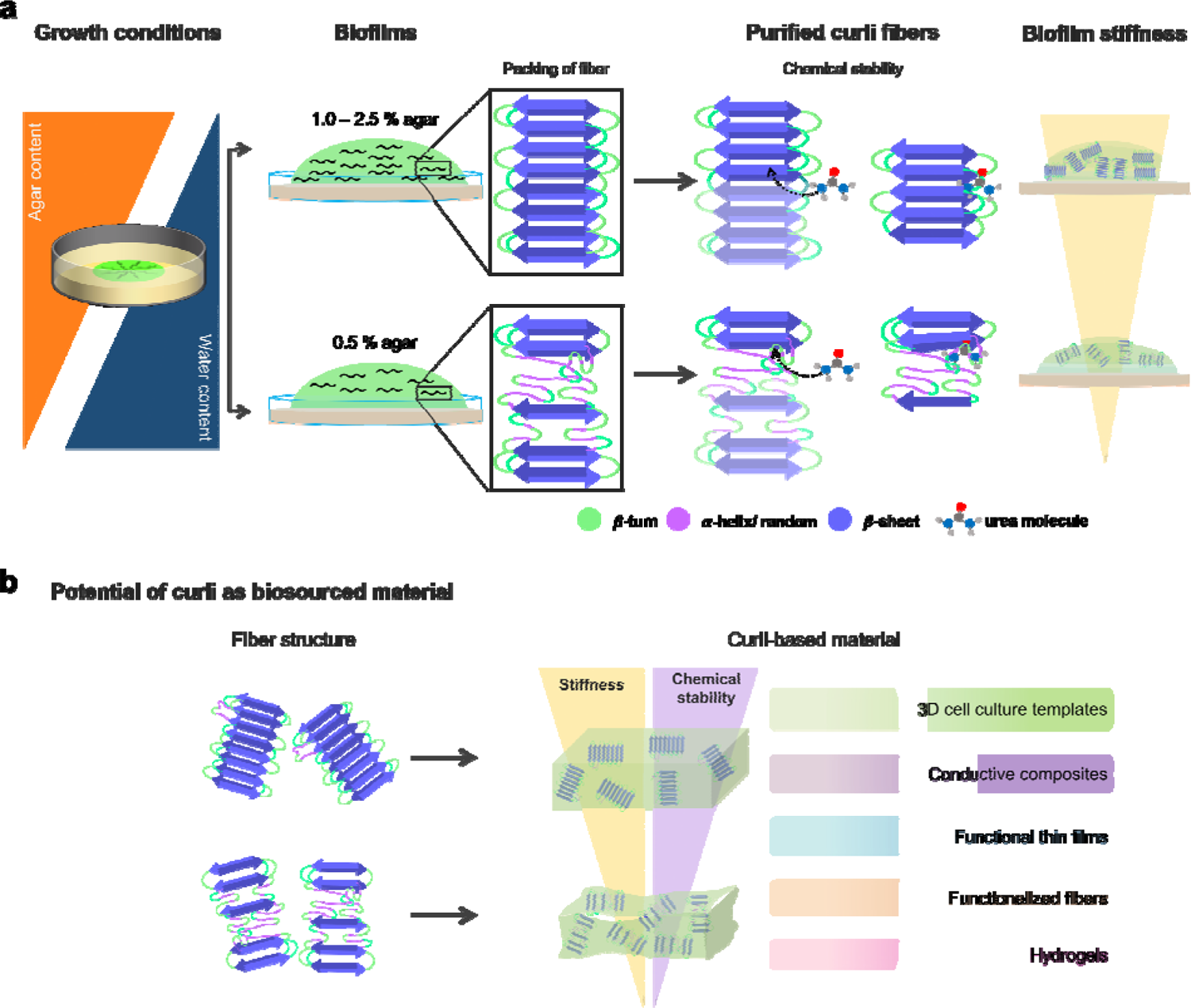
A graphical summary of the results. **(a)** By varying the water content in the substrates used for biofilm growth, we showed a dependence of the production of curli fibers. The packing of the fibers also varies as a function of the substrate water content, which in turn influences their chemical stability and the mechanical properties of the materials they constitute (e.g. a biofilm). (b) The potential of curli as a building block for bio-sourced materials^10–13,23^.

Previous studies involving different bacteria strains showed that biofilms grown on substrates with high water content (e.g. 0.5 % agar) have a less dense matrix, which is heterogeneously distributed across the biofilm thickness.^15,17^ Our results would attribute these observations to a lower production of curli fibers in these conditions, as well as to different final fiber conformation (Figure 1e, **Table 2**). By affecting the quantity and the molecular structure of curli fibers in the biofilm matrix (Figure 2c and Figure 6), the water content of the agar substrate is therefore expected to affect biofilm macroscopic features like their growth kinetics^7^ and their mechanical properties.^3,15,19^ For example, the high content of β-turns in the curli fibers purified from biofilm grown on substrates with high water content (0.5 % agar) results in a lower packing compared to the fibers obtained from dryer conditions (1.0 – 2.5 % agar) (Figure 2e). Such difference in structure could be explained by larger average spacing between strands (Figure 6a), and the resulting flexibility of the curli fibers in the matrix contribute to the lower rigidity of biofilms grown on wet substrates as measured by micro-indentation experiments (Figure 5b).^17^ It is important to note that previous studies decoupled the role of the water content and of the stiffness of the substrate using semipermeable membranes, and revealed that differences in biofilm growth behavior in such conditions are rather due to the water availability than substrates mechanics.^62^ Moreover, wet substrates were proposed to promote bacteria motility by enabling swimming, which would explain the larger biofilms obtained in such conditions, as well as their less dense matrix and softer mechanical properties.^17^ Such differences in the microenvironment of the curli fibers could in turn influence their packing and conformation.

**Table 2.** FWHH input values in cm−1 and their physically plausible ranges expected for each type of secondary structure^27,29,31^.

| Secondary structure component |  | FWHH input (cm<sup>-1</sup>) | Lower limit (cm<sup>-1</sup>) | Upper limit (cm<sup>-1</sup>) |
| --- | --- | --- | --- | --- |
| β-sheet | High wavenumber component | 9 | 8 | 11 |
|  | Low wavenumber component | 17 | 14 | 19 |
| α-helix/ random |  | 20 | 5 | 30 |
| Turns |  | 20 | 5 | 30 |

In bacterial biofilms, curli amyloid fibers influence cell properties such as their ability to withstand drying, as well as the thickness, the hydrophobicity and the rigidity of the biofilm.^15^ In this work, a more hydrophobic behavior of the curli fibers and a stiffer behavior of the biofilm follow a denser packing (Figure 3e**, 5c**)^15^ and a higher β-sheet content of the fibers (**Figure S11**). However, no obvious correlation was found between biofilm mechanics (i.e. biofilm reduced Young’s modulus) and their content of curli amyloid fibers (i.e. fiber yield). While other components such as cells, eDNA, accessory proteins, lipids and sugars are also known to influence the functional properties of the biofilm material,^15,63^ our results support that the curli fibers in the biofilm matrix also have a determining role there. Either the bacteria react to the environmental cues by forming structurally different CsgA monomers, or the fibers adopt a particular folding outside the bacteria depending on the environment.^53^ In both cases, the secondary structure of the resulting amyloid fibers would in turn contribute to the matrix and biofilm materials properties under the given conditions. For example, the biofilms grown in the extreme conditions (0.5% and 2.5% salt-free LB-agar) contained less curli and had less water uptake capacity (Figure 1f-g). These results suggest that, in addition to providing biofilms with adhesion and rigidity,^3,17^ curli fibers also promote biofilm water uptake from the surroundings, thereby contributing to their hydration capacity. This ability constitutes a significant advantage for biofilms growing at solid-air interfaces, i.e. in environments where an influx of water carries the nutrients from the substrate to the biofilm. While osmotic gradients have been proposed to be involved in this transport function of the biofilm matrix, their origin is yet not clear.^16^ Since i) bacteria are able to regulate the number and location of the curli fibers formed,^7^ and ii) less curli fibers were found in the biofilms grown in wet conditions than those grown in dryer conditions (Figure 1e), then curli production could be proposed as a way for bacteria to create these osmotic gradients in conditions where water is more difficult to reach. The discrepancy of this hypothesis with the lower quantity of curli fibers measured in biofilms grown on the driest substrates (2.5 % agar) could be explained by the extreme difficulties to reach the nutrients, which could impair the overall biofilm growth and/or matrix production.

Biofilms can be seen as natural hydrogels, in which bacteria are embedded in a cross-linked polymer matrix.^64^ This perspective contributes to the emerging trend of considering *E.coli*-grown amyloid fibers (i.e. curli) are promising building blocks for making sustainable and bio-sourced materials with interesting functional properties. As such, these biopolymers have been used to produce hydrogels,^12^ functional templates for protein immobilization,^11^ conductive composite,^13^ among others. For instance, adding curli fibers to alginate hydrogels was shown to increase their stiffness.^12^ Our work suggests that the stiffness of such hydrogels could also be adjusted by tuning the structure of the added curli fibers rather than by adjusting their amount (Figure 6a). This alternative can become interesting in situations where the mechanical properties of the materials need to be decoupled from their composition, e.g., in tissue engineering research.^65^ While genetic engineering remains the preferred approach to design curli fibers with specific characteristics for further use in functional materials,^11^ our work shows the potential of using biofilm growth conditions as an interesting alternative to tune the physico-chemical properties of curli amyloid fibers. As such, a given strain of biofilm-forming *E. coli* could yield curli fibers of different secondary structures on demands.

Overall, this work helps to understand better the effect of water availability during *E. coli* biofilm growth on the secondary structure of the curli amyloid fibers extracted from the biofilm matrix. The biophysics-based methods used to characterize curli at the molecular scale appear to be of great value to inform materials engineers about the physico-chemical properties of this promising building block. Moreover, our study reports on how these differences further impact other functional properties of the curli fibers on their own (chemical and thermal stability) or of a material where they greatly contribute (e.g. biofilm mechanical properties). Finally, the findings reported provide valuable knowledge regarding the structure – function relationship that spans biofilm scales from the molecular to the tissue levels, and give insights into the adaptation response of biofilms in specific environments. As such, this study will also contribute to uncover the full potential of biofilm matrix as building blocks to engineer living and/or bio-sourced materials, which can eventually become construction materials, sustainable plastic-like materials or scaffolds for tissue engineering (Figure 6b).^11,12,23^

### Experimental Section

#### Bacterial strain and growth

The biofilm-forming bacterial strain *E. coli* K-12 W3110 was used throughout this study. Salt-free LB-agar plates (15mm diameter) were prepared with 0.5%, 1.0%, 1.8%, or 2.5% w/v of bacteriological grade agar−agar (Roth, 2266), supplemented with 1% w/v tryptone (Roth, 8952) and 0.5% w/v yeast extract (Roth, 2363). After agar pouring, the plates were left to dry for 10 minutes with the lid open and 10 minutes with the lid partially open to avoid future condensation. Each agar plate was left to rest for 48 hours before bacteria seeding. A suspension of bacteria was prepared from a single colony and grown overnight in Luria−Bertani (LB) medium at 37°C with shaking at 250 rpm. Each plate was inoculated with arrays of 9 drops of 5μL of bacterial suspension (OD600 ∼ 0.5 after 10x dilution). After inoculation, the excess of water evaporated from the drops and left bacteria-rich dry traces of comparable sizes from 4 to 8mm diameter, depending on the growth condition. Biofilms were grown for 5 days in total (∼120h) inside an incubator at 28°C. Monitoring the relative humidity in the incubator showed that it remains around 30%RH.

#### Biofilm imaging

Three biofilms per condition were imaged with a stereomicroscope (AxioZoomV.16, Zeiss, Germany) using the tiling function of the acquisition software (Zen 2.6 Blue edition, Zeiss, Germany). To estimate the biofilm size, 3 independent biofilms were measured at their equatorial line using the Fiji software.^66^ An average was then calculated for each growth condition.

#### Biofilm water content, dry mass and water uptake

The water content and water uptake of the biofilms were determined by scraping 7 biofilms per condition from the respective agar substrates after 5 days of growth (∼120 h). Biofilms were placed in plastic weighing boats, and dried at 60°C for 3 h in an oven. Wet and dry masses (m_wet_, m_dry_) were determined before and after drying.^17^ To determine the water uptake (W_up_), we added Millipure water in excess (5 ml) to the biofilms harvested from each condition, covered them with aluminum foils to avoid evaporation and left overnight. The water excess was removed and the biofilm samples were weighed again (m_rewet_). The biofilms water content in each growth condition was estimated with **Eq. (1)**

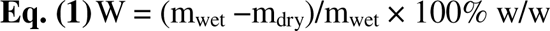

The percentage of water uptake of biofilms after rehydration (%W_up,w_) was determined with respect to biofilm initial wet mass as described in **Eq. (2)**

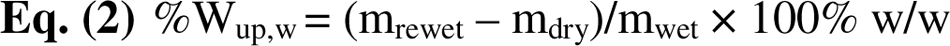

The water uptake per gram of dry biofilm (W_up,d_) was calculated with **Eq. (3)**

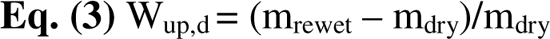

All procedures were carried out in four independent experiments.

### Curli fiber purification and quantification

Fiber purification involved a similar process as reported in previous works.^21^ Briefly, a total of 27 biofilms (**∼** 1g of biofilm material) were scraped from the surface of the substrates. Biofilms were blended five times on ice with an XENOX MHX 68500 homogenizer for 1min at 2-min intervals. The bacteria were pelleted by centrifuging two times at low speed (5000g at 4°C for 10min). A final concentration of NaCl 150 mM was added to the supernatant and the curli pelleted by centrifuging at 12.000g at 4°C for 10 minutes. The pellet was resuspended in 1mL of solution containing 10mM tris (pH 7.4) and 150mM NaCl, and incubated on ice for 30min before being centrifuged at 16.000g at 4°C for 10 minutes. This washing procedure was repeated thrice. The pellet was then resuspended in 1mL of 10mM tris solution (pH 7.4) and pelleted as described above (16.000g at 4°C for 10 minutes). The pellet was again suspended in 1mL of 10mM tris (pH 7.4) and centrifuged at 17.000g at 4°C for 10 minutes. This washing step was repeated twice. The pellet was then resuspended in 1mL of SDS 1% v/v solution and incubated for 30min. The fibers were pelleted by centrifuging at 19.000g at 4°C for 15min. The pellet was resuspended in 1mL of Milli-Q water. This washing procedure was repeated thrice. The last resuspension was done in 0.1mL of Milli-Q water supplemented with 0.02% sodium azide. The fiber suspension was stored at 4°C for later use. The protein concentration in monomeric units of the suspensions was determined by the absorbance from an aliquot incubated in 8M urea at 25°C for 2h, a treatment leading to complete dissociation of the fibrils as verified by Thioflavin T measurements.

### Transmission electron microscopy (TEM)

2μL drops of fiber suspension were adsorbed onto Formvar-coated carbon grids (200 mesh), washed with Milli-Q water, and stained with 1% (w/v) uranyl acetate. The samples were imaged in a JEOL-ARM F200 transmission electron microscope equipped with two correctors for imaging and probing. For the observations, we applied an acceleration voltage of 200 kV. The width of the purified fibers values in TEM images were measured using the scale tool of the GATAN GMS 3 software. To avoid subjectivity over 10 different images and over 10 identified fibers per field were used.

### Fluorescence confocal microscopy

Fiber samples of 5μM monomer concentration were incubated with 0.2mg/mL Thioflavin S (ThioS) for 2h at room temperature under stirring. The fibers were then washed twice by alternating centrifugation and resuspension in Milli-Q water to remove the excess of ThioS. The fibers were finally observed under a LEICA confocal microscope SP8 FALCON (Leica, Mannheim, Germany) with a water immersion 62X objective (1.2NA) under excitation at 405nm in the emission range 415-550 nm. To estimate the ThioS mean intensity in each fiber, five images were analyzed: different points in the fiber region were taken and the signal normalized according to the background signal, an average was then estimated. This procedure was performed five times in each image analyzed.

### Fluorescence spectroscopy: ThioT emission

Fiber samples were normalized to a 5µM CsgA monomer concentration (as previously described) for all fluorescence spectroscopy experiments. Corrected steady-state emission spectra were acquired with a FluoroMax®-4 spectrofluorometer (HORIBA). Spectra were recorded at 25°C using a 3-mm path cuvette (Hellma® Analytics). ThioT measurements were performed at final concentrations of 3 μM protein, 1 mM probe in Glycine buffer, pH 8.2, using λ_exc_ = 446 nm and spectral bandwidths of 10 nm.

### Circular Dichroism (CD) spectroscopy

Spectra of 5µM CsgA monomer concentration of fiber solution (as previously described) in Milli-Q water were recorded with a Chirascan CD spectrometer (Applied Photophysics, Leatherhead, Surrey, UK). A quartz cuvette with 1 mm path length (Hellma, Müllheim, Germany) was used. Spectra are acquired between 190 nm and 250 nm wavelengths, with 1 nm step size, 1 nm band-width and 0.7 s integration time per point. Milli-Q water was used to define the measurement background, which was automatically subtracted during acquisition. The experiments were repeated thrice for each condition. Each measurement was an average of three scans.

### Attenuated total reflectance Fourier transform infrared spectroscopy (ATR-FTIR)

IR spectra were acquired on a spectrophotometer (Vertex 70v, Bruker Optik GmbH, Germany) equipped with a single reflection diamond reflectance accessory continuously purged with dry air to reduce water vapor distortions in the spectra. Fibers in Milli-Q water samples (∼10μL) were spread on a diamond crystal surface, dried under N_2_ flow to obtain the protein spectra. A total of 64 accumulations were recorded at 25°C using a nominal resolution of 4cm^−1^.

Spectra were processed using Kinetic software developed by Dr. Erik Goormaghtigh at the Structure and Function of Membrane Biology Laboratory, Université Libre de Bruxelles, Brussels, Belgium. After subtraction of water vapor and side chain contributions, the spectra were baseline corrected and area normalized between 1700 and 1600cm^−1^ (Figure S2). For a better visualization of the overlapping components arising from the distinct structural elements, the spectra were deconvoluted using Lorentzian deconvolution factor with a full width at the half maximum (FWHM) of 20 cm^−1^ and a Gaussian apodization factor with a FWHM of 30 cm^−1^ to achieve a line narrowing factor K = 1.5.^31^ Second derivative was performed on the Fourier self-deconvoluted spectra for band assignment. The bands identified by both procedures were used as initial parameters for a least square iterative curve fitting of the original IR band (K = 1) in the amide I’ region, using mixed Gaussian/Lorentzian bands. Peak positions of each identified individual component were constrained within ±2 cm^−1^ of the initial value.

### Fluorescence spectroscopy: 1,8-ANS emission

Corrected steady-state emission spectra were acquired with a FluoroMax®-4 spectrofluorometer (HORIBA). Spectra were recorded at 25°C using a 3-mm path cuvette (Hellma® Analytics). 1-anilino-nafthalene-8-sulfonate (1,8-ANS) emission spectra were obtained after 30 min incubation at room temperature of mixture containing 5 μM fiber concentration in monomeric units and 100 μM dye in the corresponding buffer, using λ_exc_ = 370 nm and bandwidth of 5 nm.

### Phasor analysis

Spectral phasor plots were used to visualize 1,8-ANS spectral shifts. The phasor plots are 2D scatter graphs, where the axes are the real (G) and imaginary (S) components of the Fourier transform of the fluorescence spectra. This transformation offers a powerful, model free, graphic method to characterize spectral^36^ and lifetime information.^67^ A detailed analysis can be found elsewhere.^41,68,69^ In this work, ANS spectra can be transformed using the following for x and y coordinates:

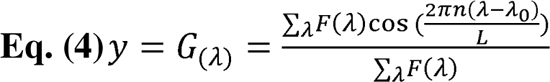

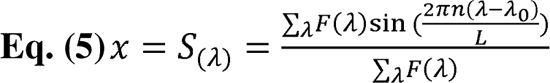

where F(λ) are the fluorescence intensity values, λ_0_ is the initial wavelength of the spectrum (λ_0_ = 400 nm), L is the length of the spectrum and n is the harmonic value. The values L=300 (from 400 to 700 nm) and n=1 were used for phasor calculations. The angular position of the phasor in the plot is related to the center of mass, while the radial position depends on the full width at half maximum of the spectrum. Spectral phasor plots were constructed using Originlab data analysis software.

### Fluorescence Lifetime Imaging Microscopy (FLIM)

Two-photon excitation of Nile red was performed at 860 nm with a pulsed Ti:Sapphire laser with 80Hz repetition rate (Spectra-Physics Mai Tai, Mountain View, CA). The image size was 512×512 pixels, the pixel size was 75 nm and the detection range were 600 to 728 nm. FLIM calibration of the system was performed by measuring the known lifetime of the fluorophore Coumarin 6 in ethanol.^41^ For these experiments, 5 μM fibers where placed in clean glass and 5 μM of NR was added.

The FLIM phasor analysis allows the transformation of the fluorescence signal from each pixel in the image to a point in the phasor plot. FLIM data were processed using SimFCS, an open source software developed at the Laboratory of Fluorescence Dynamics, Irvine, California (available at http://www.lfd.uci.edu). Further details of FLIM phasor analysis are provided in S4.

### Fluorescence spectroscopy: intrinsic fiber fluorescence and quenching experiments

Fiber samples were normalized to a 5µM monomer concentration for all fluorescence spectroscopy experiments. Corrected steady-state emission spectra were acquired with a FluoroMax®-4 spectrofluorometer (HORIBA). Spectra were recorded at 25°C using a 3-mm path cuvette (Hellma® Analytics). Intrinsic fluorescence spectra (5 μM protein) were acquired using λ_exc_= 280 nm and 5/5 nm slit bandwidths. For quenching experiments, increasing volumes of 5 M acrylamide or potassium iodide (KI) were added to a 4 μM fiber suspension, 4 μM N-Acetyl-L-tryptophanamide (NATA, Sigma-Aldrich) solution or buffer alone until reaching a maximum quencher concentration of 0.37 M. Spectra were recorded after 5 min incubation using λ_exc_ = 280 nm. Blank substracted spectra were corrected for dilution and inner filter effects by multiplication with a factor of 10(A_exc_+A_em_)/2, thereby compensating for the absorption of KI and acrylamide at the fluorescence excitation and emission wavelength. Quenching curves were fit by linear regression with the Stern-Volmer equation (**Eq. (3)**)

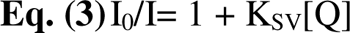

where I_0_ and I are the fluorescence intensities in the absence and in the presence of quencher, respectively, at the concentration [Q] and K_SV_ is the dynamic quenching constant.

### Thermal stability assay

5μM fiber samples were incubated at 25, 30, 40, 50, 60, 70 and 80°C for 3 min before their measurement with ATR-FTIR spectroscopy. An extra measurement was done by taking the fibers from 80°C to a bath water at 25°C and incubate the samples there for 3 min. Data were acquired as described above.

### Chemical stability assay

To test the chemical stability of the protein fibers, 5 μM samples were prepared by incubation of increasing urea concentration (0 – 8 M) and left for 2 h at room temperature to ensure equilibrium. Thioflavin T (ThioT) was then added in a final concentration of 1 mM and fluorescence emission spectra of the samples were acquired under excitation at λ_exc_ = 446 nm and spectral bandwidth of 5 nm. Emission of the ThioT in urea was done and no significant signal was detected.

The IC50 value was estimated by fitting the data to a linear regression (Y = a * X + b) and then calculating IC50 = (0.5 - b)/a.

#### Micro-indentation on biofilms

The micro-indentation experiments were carried out as described in Ziege et al.^17^ Briefly, *E. coli* W3110 biofilms were grown for 5 days on their respective agar substrate and 3 to 4 of them were tested per condition. Ten measurements were performed in the central region of each biofilm. The distance between two measurement points was at least 250 μm in x and y directions and the depth of the indentation was between 10 and 30 µm, i.e. much less than biofilm the thickness (∼100µm).^70^ A TI 950 Triboindenter (Hysitron Inc.) was used to determine the load– displacement curves after calibration of the instrument in air. Loading rates ranged from 20 to 30 μm/s, which translates to loading and unloading times of 10 s. The loading portion of all curves were fitted with a Hertzian contact model over and indentation range of 0 to 10µm to obtain the reduced Young’s modulus E_r_.

### Statistical analysis

For each experiment, 3 to 4 fiber solutions were used, where each solution came from different fiber purification batches. For each purification, 27 biofilms were cultured in each of the 4 growth conditions tested (i.e. on salt-free LB-agar plate containing 0.5%, 1.0%, 1.8% and 2.5% agar respectively). For each batch of biofilm culture, the different samples of fibers obtained for the 4 conditions were treated simultaneously (or in consecutive days) to avoid variability due to unavoidable slight variations in the implementation of the protocols (e.g. temperature and humidity in the laboratory during agar preparation and/or biofilm seeding).

For statistical analysis, a Shapiro Wilk test was used to check for data normality. For data with no normal distribution, Kruskal-Wallis non-parametric test was performed. For data with normal distribution, a One-way ANOVA test was carried out. Mechanical properties data was analyzed using a Mann-Whitney U test. Unless otherwise stated in the caption, Dunn’s post-test for multiple comparisons were done with respect to the 1.8 % salt-free LB-agar condition, considered as the standard seeding condition. Details of each test are described in the legend of the figures.

## Supporting Information

The following files are available free of charge:

- Table S1: Composition of salt-free LB agar subsrates
- Figure S1: Characterization of salt-free LB agar substrates
- Figure S2: Biofilm dry mass
- Figure S3: Purity of curli fibers
- Figure S4: ζ-potential of purified curli fibers
- Figure S5: Amide I’ spectra of curli fibers purified from *E. coli* biofilms
- Figure S6: Spectral phasor plot analysis
- Figure S7: FLIM phasor plot
- Figure S8: Nile red FLIM analysis
- Figure S9: Raw spectra of the quenching experiments
- Table S2: Stern Volmer constants of quenching experiments
- Figure S10: Complete fiber thermal denaturation ramp (ATR-FTIR analysis)
- Figure S11: Micro-indentation curves of *E. coli* W3110 biofilms
- Figure S12: Fiber structure/ fiber function relationship

## Data Availability Statement

The data that support the findings of this study are available from the corresponding authors upon reasonable request.

## AUTHOR INFORMATION

### Author Contributions

The manuscript was written through contributions of all authors. All authors have given approval to the final version of the manuscript. The authors declare no competing interests.

## ACKNOWLEDGMENT

M.S. acknowledges support from the Max Planck Queensland Centre on the Materials Science for Extracellular Matrices and A.M acknowledges support from the Alexander von Humboldt foundation. The authors also thank Christine Pilz-Allen for her technical support in the laboratories, Peter Werner and Heike Runge for their help in doing the transmission electronic microscopy experiments, and Peter Fratzl and Ricardo Ziege for the useful discussions throughout this study. The authors are also grateful to Regine Hengge (HU Berlin) for providing the *E. coli* strain W3110 and to Eric Goormaghtigh from the SFMB group at the Université Libre de Bruxelles for providing the Kinetics Software.

## REFERENCES

1. Jeffries, J., Fuller, G. G. & Cegelski, L. Unraveling Escherichia coli’s Cloak: Identification of Phosphoethanolamine Cellulose, Its Functions, and Applications. Microbiol. Insights 12, 117863611986523 (2019).

2. Jeffries, J. et al. Variation in the ratio of curli and phosphoethanolamine cellulose associated with biofilm architecture and properties. Biopolymers 112, 1–11 (2021).

3. Serra, D. O., Richter, A. M. & Hengge, R. Cellulose as an architectural element in spatially structured escherichia coli biofilms. J. Bacteriol. 195, 5540–5554 (2013).

4. Akbey, Ü. & Andreasen, M. Functional amyloids from bacterial biofilms – structural properties and interaction partners. Chem. Sci. 13, 6457–6477 (2022).

5. Dueholm, M. S. et al. Fibrillation of the major curli subunit CsgA under a wide range of conditions implies a robust design of aggregation. Biochemistry 50, 8281–8290 (2011).

6. Evans, M. L. & Chapman, M. R. Curli biogenesis: Order out of disorder. Biochim. Biophys. Acta - Mol. Cell Res. 1843, 1551–1558 (2014).

7. Andreasen, M. et al. Physical determinants of amyloid assembly in biofilm formation. MBio 10, 1–12 (2019).

8. Romero, D., Aguilar, C., Losick, R. & Kolter, R. Amyloid fibers provide structural integrity to Bacillus subtilis biofilms. Proc. Natl. Acad. Sci. U. S. A. 107, 2230–2234 (2010).

9. Knowles, T. P. J. & Mezzenga, R. Amyloid fibrils as building blocks for natural and artificial functional materials. Adv. Mater. 28, 6546–6561 (2016).

10. Dorval Courchesne, N. M., Duraj-Thatte, A., Tay, P. K. R., Nguyen, P. Q. & Joshi, N. S. Scalable Production of Genetically Engineered Nanofibrous Macroscopic Materials via Filtration. ACS Biomater. Sci. Eng. 3, 733–741 (2017).

11. Nguyen, P. Q., Botyanszki, Z., Tay, P. K. R. & Joshi, N. S. Programmable biofilm-based materials from engineered curli nanofibres. Nat. Commun. 5, 1–10 (2014).

12. Axpe, E. et al. Fabrication of Amyloid Curli Fibers-Alginate Nanocomposite Hydrogels with Enhanced Stiffness. ACS Biomater. Sci. Eng. 4, 2100–2105 (2018).

13. Huyer, C. et al. Fabrication of Curli Fiber-PEDOT:PSS Biomaterials with Tunable Self-Healing, Mechanical, and Electrical Properties. ACS Biomater. Sci. Eng. (2021) doi:10.1021/acsbiomaterials.1c01180.

14. Chen, D. et al. Characteristics and influencing factors of amyloid fibers in S. mutans biofilm. AMB Express 9, 1–9 (2019).

15. Zeng, G. et al. Functional bacterial amyloid increases Pseudomonas biofilm hydrophobicity and stiffness. Front. Microbiol. 6, 1–14 (2015).

16. Trinschek, S., John, K., Lecuyer, S. & Thiele, U. Continuous versus Arrested Spreading of Biofilms at Solid-Gas Interfaces: The Role of Surface Forces. Phys. Rev. Lett. 119, 1–5 (2017).

17. Ziege, R. et al. Adaptation of Escherichia coli Biofilm Growth, Morphology, and Mechanical Properties to Substrate Water Content. ACS Biomater. Sci. Eng. 7, 5315–5325 (2021).

18. Ryzhkov, N. V., Nikitina, A. A., Fratzl, P., Bidan, C. M. & Skorb, E. V. Polyelectrolyte Substrate Coating for Controlling Biofilm Growth at Solid–Air Interface. Adv. Mater. Interfaces 8, (2021).

19. Lembré, P., Di Martino, P. & Vendrely, C. Amyloid peptides derived from CsgA and FapC modify the viscoelastic properties of biofilm model matrices. Biofouling 30, 415–426 (2014).

20. Thongsomboon, W. et al. Phosphoethanolamine cellulose: A naturally produced chemically modified cellulose. Science (80-.). 359, 334–338 (2018).

21. Chapman, M. R. et al. Role of Escherichia coli curli operons in directing amyloid fiber formation. Science (80-.). 295, 851–855 (2002).

22. Chiti, F. & Dobson, C. M. Protein Misfolding, Functional Amyloid, and Human Disease. Annu. Rev. Biochem. 75, 333–366 (2006).

23. Duraj-Thatte, A. M. et al. Water-processable, biodegradable and coatable aquaplastic from engineered biofilms. Nat. Chem. Biol. 17, 732–738 (2021).

24. Sun, A., Nguyen, X. V. & Bing, G. Comparative analysis of an improved thioflavin-S stain, Gallyas silver stain, and immunohistochemistry for neurofibrillary tangle demonstration on the same sections. J. Histochem. Cytochem. 50, 463–472 (2002).

25. Krebs, M. R. H., Bromley, E. H. C. & Donald, A. M. The binding of thioflavin-T to amyloid fibrils: Localisation and implications. J. Struct. Biol. 149, 30–37 (2005).

26. Nagaraj, M. et al. Predicted Loop Regions Promote Aggregation: A Study of Amyloidogenic Domains in the Functional Amyloid FapC. J. Mol. Biol. (2020) doi:10.1016/j.jmb.2020.01.044.

27. Siri, M., Herrera, M., Moyano, A. J. & Celej, M. S. Influence of the macromolecular crowder alginate in the fibrillar organization of the functional amyloid FapC from Pseudomonas aeruginosa. Arch. Biochem. Biophys. 713, 109062 (2021).

28. Arad, E., Green, H., Jelinek, R. & Rapaport, H. Revisiting thioflavin T (ThT) fluorescence as a marker of protein fibrillation – The prominent role of electrostatic interactions. J. Colloid Interface Sci. 573, 87–95 (2020).

29. Sarroukh, R., Goormaghtigh, E., Ruysschaert, J. M. & Raussens, V. ATR-FTIR: A ‘rejuvenated’ tool to investigate amyloid proteins. Biochim. Biophys. Acta - Biomembr. 1828, 2328–2338 (2013).

30. Moran, S. D. & Zanni, M. T. How to get insight into amyloid structure and formation from infrared spectroscopy. J. Phys. Chem. Lett. 5, 1984–1993 (2014).

31. Goormaghtigh, E., Cabiaux, V. & Ruysschaert, J. M. Secondary structure and dosage of soluble and membrane proteins by attenuated total reflection Fourier[transform infrared spectroscopy on hydrated films. Eur. J. Biochem. 193, 409–420 (1990).

32. Barnhart, M. M. & Chapman, M. R. Curli biogenesis and function. Annu. Rev. Microbiol. 60, 131–147 (2006).

33. Hammarström, P., Lindgren, M. & Nilsson, K. P. R. Fluorescence Spectroscopy as a Tool to Characterize Amyloid Oligomers and Fibrils. in Amyloid Fibrils and Prefibrillar Aggregates 211–243 (2013). 10.1002/9783527654185.ch11.

34. Sarell, C. J. et al. Expanding the Repertoire of Amyloid Polymorphs by Co-polymerization of Related Protein Precursors*. J. Biol. Chem. 288, 7327–7337 (2013).

35. Younan, N. D. & Viles, J. H. A Comparison of Three Fluorophores for the Detection of Amyloid Fibers and Prefibrillar Oligomeric Assemblies. ThT (Thioflavin T); ANS (1-Anilinonaphthalene-8-sulfonic Acid); and bisANS (4,4′-Dianilino-1,1′-binaphthyl-5,5′-disulfonic Acid). Biochemistry 54, 4297–4306 (2015).

36. Fereidouni, F., Bader, A. N. & Gerritsen, H. C. Spectral phasor analysis allows rapid and reliable unmixing of fluorescence microscopy spectral images. Opt. Express 20, 12729 (2012).

37. Malacrida, L. & Gratton, E. LAURDAN fluorescence and phasor plots reveal the effects of a H2O2 bolus in NIH-3T3 fibroblast membranes dynamics and hydration. Free Radic. Biol. Med. 128, 144–156 (2018).

38. Hawe, A., Sutter, M. & Jiskoot, W. Extrinsic fluorescent dyes as tools for protein characterization. Pharm. Res. 25, 1487–1499 (2008).

39. Scheibel, T. & Serpell, L. Physical Methods for Studies of Fiber Formation and Structure. Protein Folding Handbook vol. 1 (2008).

40. Mishra, R., Sjölander, D. & Hammarström, P. Spectroscopic characterization of diverse amyloid fibrils in vitro by the fluorescent dye Nile red. Mol. Biosyst. 7, 1232–1240 (2011).

41. Digman, M. A., Caiolfa, V. R., Zamai, M. & Gratton, E. The phasor approach to fluorescence lifetime imaging analysis. Biophys. J. 94, 14–16 (2008).

42. Sancataldo, G., Anselmo, S. & Vetri, V. Phasor-FLIM analysis of Thioflavin T self-quenching in Concanavalin amyloid fibrils. Microsc. Res. Tech. 83, 811–816 (2020).

43. Chung, C. W. et al. Label-Free Characterization of Amyloids and Alpha-Synuclein Polymorphs by Exploiting Their Intrinsic Fluorescence Property. Anal. Chem. 94, 5367–5374 (2022).

44. Levitt, J. A., Chung, P.-H. & Suhling, K. Spectrally resolved fluorescence lifetime imaging of Nile red for measurements of intracellular polarity. J. Biomed. Opt. 20, 096002 (2015).

45. Golini, C. M., Williams, B. W. & Foresman, J. B. Further Solvatochromic, Thermochromic, and Theoretical Studies on Nile Red. J. Fluoresc. 8, 395–404 (1998).

46. Shu, Q. et al. The E. coli CsgB nucleator of curli assembles to β-sheet oligomers that alter the CsgA fibrillization mechanism. Proc. Natl. Acad. Sci. U. S. A. 109, 6502–6507 (2012).

47. Ladokhin, A. S. Fluorescence Spectroscopy in Peptide and Protein Analysis. *Encycl*. Anal. Chem. 5762– 5779 (2000) doi:10.1002/9780470027318.a1611.

48. Jumper, J. et al. Highly accurate protein structure prediction with AlphaFold. Nature 596, 583–589 (2021).

49. Eftink, M. R. & Ghiron, C. A. Fluorescence quenching studies with proteins. Anal. Biochem. 114, 199–227 (1981).

50. Lakowicz, J. R. Principles of fluorescence spectroscopy, 3rd Principles of fluorescence spectroscopy, Springer, New York, USA, 3rd edn, 2006. Principles of fluorescence spectroscopy, Springer, New York, USA, 3rd edn, 2006. (2006). doi:10.1007/978-0-387-46312-4.

51. Jiskoot, W., Hlady, V., Naleway, J. J. & Herron, J. N. Application of fluorescence spectroscopy for determining the structure and function of proteins. Pharm. Biotechnol. 7, 1–63 (1995).

52. Stevenson, S. G. & Preston, K. R. Intrinsic fluorescence and quenching studies of gluten proteins. Cereal Chem. 71, 155–158 (1994).

53. John, T., Martin, L. L. & Abel, B. Peptide Self[Assembly into Amyloid Fibrils at Hard and Soft Interfaces—From Corona Formation to Membrane Activity. Macromol. Biosci. 2200576 (2023) doi:10.1002/mabi.202200576.

54. Ball, P. Water as an active constituent in cell biology. Chem. Rev. 108, 74–108 (2008).

55. Thirumalai, D., Reddy, G. & Straub, J. E. Role of water in protein aggregation and amyloid polymorphism. Acc. Chem. Res. 45, 83–92 (2012).

56. Gohlke, J. R. the most popular fluorescence probes in use at this time. Weber. (1972).

57. Matulis, D. & Lovrien, R. 1-Anilino-8-Naphthalene Sulfonate Anion-Protein Binding Depends Primarily on Ion Pair Formation. Biophys. J. 74, 422–429 (1998).

58. Sackett, D. L. & Wolff, J. Nile red as a polarity-sensitive fluorescent probe of hydrophobic protein surfaces. Anal. Biochem. 167, 228–234 (1987).

59. Dandurand, J. et al. Conformational and thermal characterization of a synthetic peptidic fragment inspired from human tropoelastin: Signature of the amyloid fibers. Pathol. Biol. 62, 100–107 (2014).

60. Wang, X. & Chapman, M. R. Curli provide the template for understanding controlled amyloid propagation. Prion 2, 57–60 (2008).

61. Zou, Q., Habermann-Rottinghaus, S. M. & Murphy, K. P. Urea effects on protein stability: Hydrogen bonding and the hydrophobic effect. Proteins Struct. Funct. Genet. 31, 107–115 (1998).

62. Yan, J., Nadell, C. D., Stone, H. A., Wingreen, N. S. & Bassler, B. L. Extracellular-matrix-mediated osmotic pressure drives Vibrio cholerae biofilm expansion and cheater exclusion. Nat. Commun. 8, (2017).

63. Zhang, Q. et al. Mechanical Resilience of Biofilms toward Environmental Perturbations Mediated by Extracellular Matrix. Adv. Funct. Mater. 32, (2022).

64. Wilking, J. N., Angelini, T. E., Seminara, A., Brenner, M. P. & Weitz, D. A. Biofilms as complex fluids. 36, 385–392 (2011).

65. Wang, X. et al. Decoupling polymer properties to elucidate mechanisms governing cell behavior. Tissue Eng. Part B. Rev. 18, 396–404 (2012).

66. Schindelin, J., et al. Fiji: An open-source platform for biological-image analysis. Nat. Methods 9, 676–682 (2012).

67. Jameson, D. M., Gratton, E. & Hall, R. D. The measurement and analysis of heterogeneous emissions by multifrequency phase and modulation fluorometry. Appl. Spectrosc. Rev. 20, 55–106 (1984).

68. Ranjit, S., Malacrida, L., Jameson, D. M. & Gratton, E. Fit-free analysis of fluorescence lifetime imaging data using the phasor approach. Nat. Protoc. 13, 1979–2004 (2018).

69. Malacrida, L., Ranjit, S., Jameson, D. M. & Gratton, E. The Phasor Plot: A Universal Circle to Advance Fluorescence Lifetime Analysis and Interpretation. Annu. Rev. Biophys. 50, 575–593 (2021).

70. Serra, D. O., Klauck, G. & Hengge, R. Vertical stratification of matrix production is essential for physical integrity and architecture of macrocolony biofilms of Escherichia coli. Environ. Microbiol. 17, 5073–5088 (2015).

