## Supporting document for "Curli amyloid fibers in *Escherichia coli* biofilms: the influence of water availability on their structure and functional properties"

### Supporting Information

| Characterization of the salt-free LB-agar substrate | Table S1 |
| --- | --- |
|  | Figure S1 |
| Biofilm dry mass | Figure S2 |
| Purity of curli fiber | Figure S3 |
| ζ-potential of purified curli fibers | Figure S4 |
| FT-IR spectra of the purified curli fibers | Figure S5 |
| Spectral phasor plots | Figure S6 |
| FLIM phasors plots | Figure S7 |
| Nile red FLIM analysis | Figure S8 |
| Raw spectra of the quenching experiments and Ksv values | Figure S9 |
|  | Table S2 |
| Thermal stability | Figure S10 |
| Micro-indentation of *E. coli* W3110 biofilms | Figure S11 |
| Fiber structure/ fiber function relationship | Figure S12 |

#### 1. Characterization of the salt-free LB-agar substrate

1.1. **Experimental section**
     1.1.1. Salt-free LB-agar substrate plate preparation

Salt-free LB-agar plates (15mm diameter) were prepared with 0.5%, 1.0%, 1.8%, or 2.5% w/v of bacteriological grade agar−agar (Roth, 2266), supplemented with 1% w/v tryptone (Roth, 8952) and 0.5% w/v yeast extract (Roth, 2363) (Table S1). After agar pouring, the plates were left to dry for 10 minutes with the lid off and 10 minutes with the lid partially off to avoid future condensation. Each agar plate was left to rest for 48 hours before bacteria seeding to ensure the correct evaporation of possible water excess. After 5 days at 28°C, the agar plates were ready for characterization.

**Table S1** Nominal and effective water contents respectively calculated and measured for various nominal agar concentrations supplemented with 1.5 w/v% nutrients^1^.

| **Nominal agar concentration (w/v%)** | **0.5** | **1.0** | **1.8** | **2.5** |
| --- | --- | --- | --- | --- |
| **Nutrient concentration (w/v%)** | 1.5 | 1.5 | 1.5 | 1.5 |
| **Nominal water content (w/w%)** | 98.0 | 97.5 | 96.8 | 96.0 |

1.1.2. Salt-free LB-agar water content

The water content in each salt-free LB-agar substrate was determined by placing ~2 g of agar in plastic weighing boats, and dried at 60°C for 3 h in an oven. Wet and dry masses (m_wet_, m_dry_) were determined before and after drying.^1^ Their water content in each growth condition was estimated with **Eq. (1)**

**Eq. (1)**	W = (m_wet_ −m_dry_)/m_wet_ × 100% w/w

The experiment was repeated three times on independent samples.

1.1.3. Thermal Gravimetric Analysis (TGA)

TGA measurement has been performed using a Thermo Microbalance TG 209 F1 Libra (Netzsch, Selb, Germany). A platinum crucible was used for the measurement of 10 ± 5mg of samples in a Nitrogen flow of 20ml/min and a purge flow of 20ml/min (Oxygenflow 10ml/min) at a heating rate of 10K/min from 25 to 600 °C. Proteus (6.1.0) and Quadstar (7.03, MID modus) software package was used for data acquisition and analysis. The experiment was repeated three times on independent samples.

1.1.4. Micro-indentation of salt-free LB-agar substrates

The micro-indentation experiments were carried out as described in Ziege et al.^1^ After salt-free LB-agar substrate preparation, 2-3 agar plates were used for micro-indentation. Seven measurements were performed in each plate. The distance between two measurement points was at least 250 μm in x and y directions and the depth of the indentation was between 10 and 30 µm. A TI 950 Triboindenter (Hysitron Inc.) was used to determine the load–displacement curves after calibration of the instrument in air. Loading rates ranged from 20 to 30 μm/s, which translates to loading and unloading times of 10 s. The loading portion of all curves were fitted with a Hertzian contact model over and indentation range of 0 to 10µm to obtain the reduced Young’s modulus E_r_.

1.2. **Results**

Salt-free LB-agar substrates were prepared in order to have nominal water content as described in **Table S1**. A detailed characterization of the substrates was carried out in order to study the effective water content in each of the salt-free LB-agar substrates and their mechanical properties (**Figure S1**). We analyzed the water content of the salt-free LB-agar substrates by dehydration (**Figure S1a**). “Wet substrates” (0.5 % agar content) have 97.67 % of water, while “dry substrates” (2.5 % agar content) have 94.07 % of water. The intermediate conditions 1.0 and 1.8 % agar content have 97.07 and 96.17 % water content, respectively.

To characterize the water variations in the salt-free LB-agar substrates, TGA experiments were carried out. The weight loss curves obtained indicate the fraction of free (or bulk) water in the sample (**Figure S1b**).^2^ We studied the behavior of the salt-free LB-agar substrates from 25 up to 600 °C. Throughout the experiment, we were able to identify two main weight losses (**Figure 1b, inset**). The first one occurs around 150 °C for all samples and corresponds to the release of free water. The second one, between 200 and 250 °C, correlates with the rupture of the polymer network and the difference in water loss is due to the density of the polymer network.^2^ Results in these experiments suggest that the substrate with higher agar concentration have less free water (**Figure S1c**).

These differences are expected to yield differences in mechanical properties of the salt-free LB-agar substrates, namely, their stiffness (**Figure S1d**). We observed that although the differences in water content between the extreme conditions is limited (about 3 %w/w), their effect in the stiffness of the substrates is pronounced. The salt-free LB-agar substrates with the lowest water content (0.5 % agar) have a stiffness of 10.35 kPa ± 1.35, while the salt-free agar substrates with the highest water content (2.5 % agar) have a stiffness of 448.2 kPa ± 40.12.

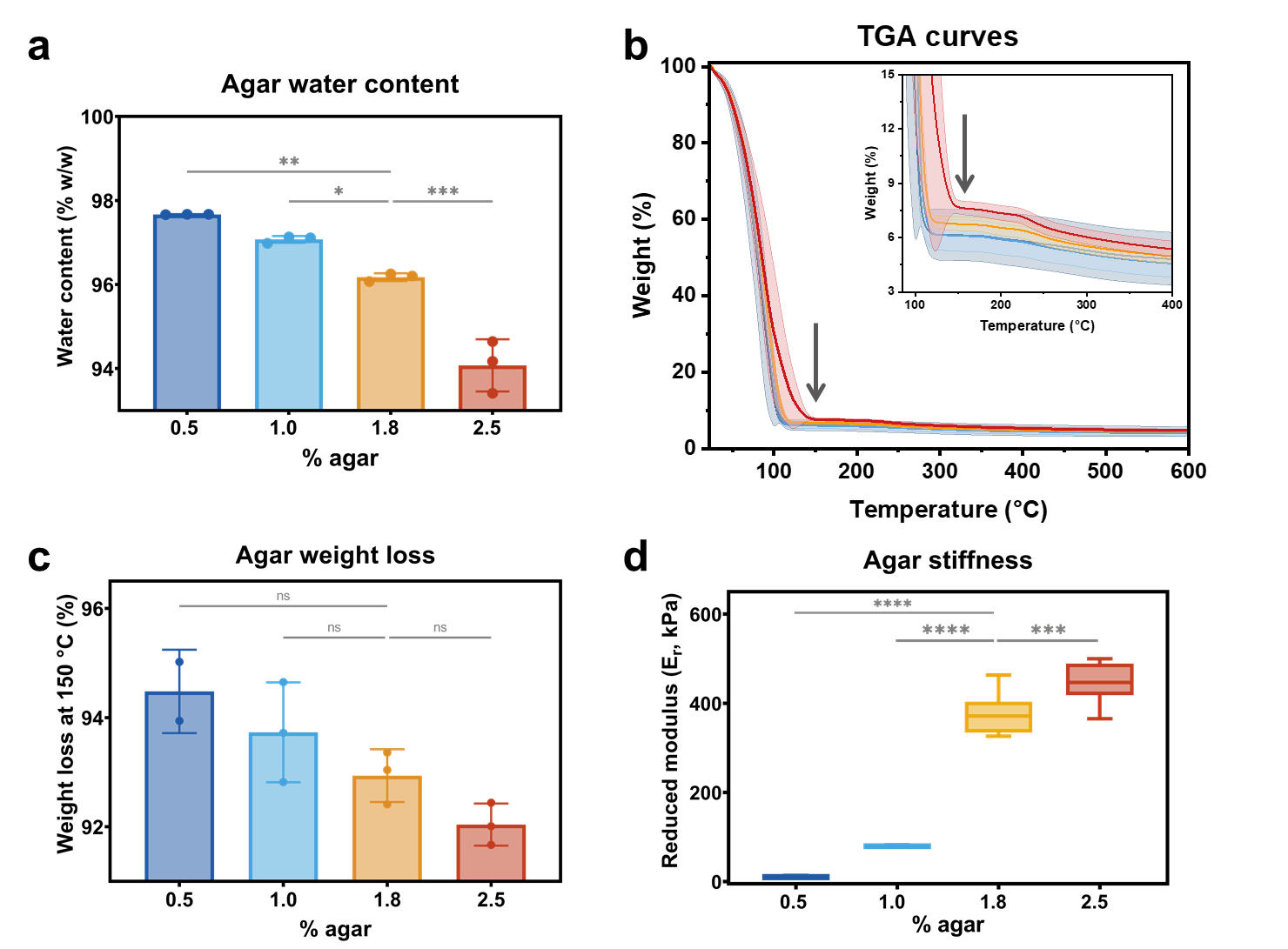

Figure S1 Agar substrate properties. (a) Water content of the different salt-free LB-agar substrates (% w/w). N=3 independent samples per condition. For statistical analysis On-way ANOVA was used (p<0.001, *** | p<0.01, ** | p<0.05, * | ns = non-significant) with a Dunnett’s post-test for multiple comparisons, comparing all conditions to the 1.8 % agar substrate. (b) Average Thermal Gravimetric Analysis (TGA) curves of the different salt-free LB-agar substrates. Inset: zoom of the curves between 85 and 400 °C. N=2-3 independent samples per condition. (c) Percentage of weight loss representing water loss at 150 °C for each substrate. For statistical analysis On-way ANOVA was used (p>0.05, ns = non-significant) with a Dunnett’s post-test for multiple comparisons, comparing all conditions to the 1.8 % agar substrate. (d) Reduced Young’s modulus of the different salt-free LB-agar substrates obtained by micro-indentation. n=14 indentation curves acquired on N=2 to 3 substrates per condition. For statistical analysis a Mann-Whitney test (p<0.0001, **** | p<0.001, ***) was carried out comparing each condition to the standard one (1.8 % agar content).

#### 2. Biofilm dry mass

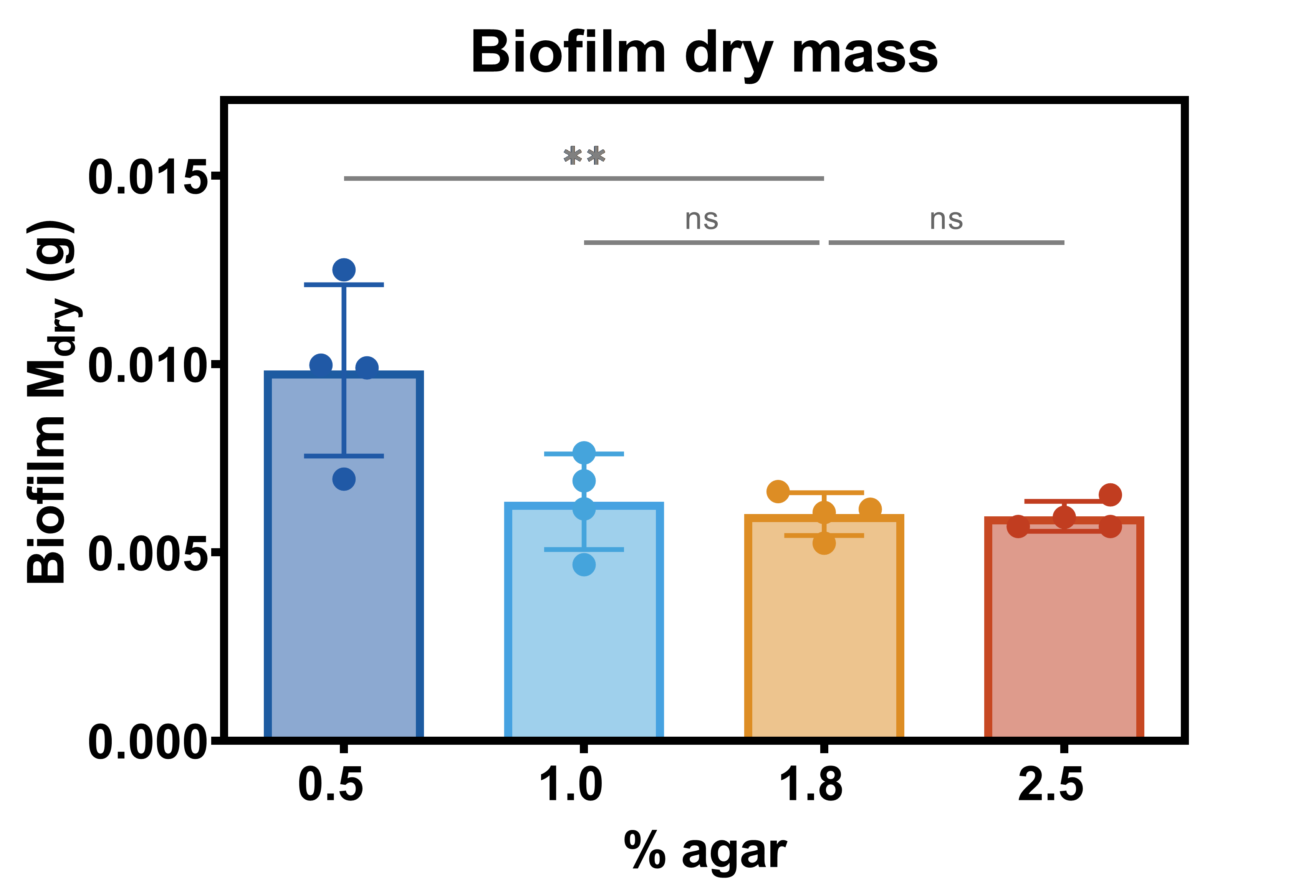

Figure S2 Biofilm dry mass. The biofilm dry mass derived from the water content in biofilms of each growth condition. N= 4 independent experiments. For statistical analysis On-way ANOVA was used (p<0.01, ** | ns = non-significant) with a Dunnett’s post-test for multiple comparisons, comparing all conditions to the 1.8 % agar substrate.

#### 3. Purity of curli fibers

We checked for chemical components or impurities 1) during and 2) after the purification of the curli fibers. We first collected 10 µL of each sample after the incubation on ice with 150 mM NaCl 10 mM Tris-HCl buffer pH 7.4, and then 10 µL of the final purified product after incubation at 37 °C in 1% SDS. We ran 12% acrylamide SDS-PAGE gels of these two samples in search of protein impurities (**Figure S3a**). The FT-IR spectra of purified curli fibers were also compared against spectra from purified *E. coli* modified-cellulose (phosphoethanolamine-cellulose) and a mixture of both components (**Figure S3b**). We confirmed the presence of purified curli fibers by TEM imaging (**Figure S3c**). This last experiment also served to see whether we had remains of cell envelope components in our samples. **In all cases, we did not fine impurities or other cell envelop components in the purified curli fibers.**

After assessing the purity of our sample, we studied whether the purification process alters the structure of curli fibers in a significant manner (**Figure S3d-f**). For this, we repeated the two main steps of the purification process on the freshly purified curli fibers: 1) an incubation of the fibers on ice with 150 mM NaCl 10 mM Tris-HCl buffer at pH 7.4, followed by 2) an incubation at 37 °C in 1% SDS. The ATR-FTIR and CD spectroscopy data collected after each of these steps suggest no significant changes in the structure of the curli fibers during these purification steps.

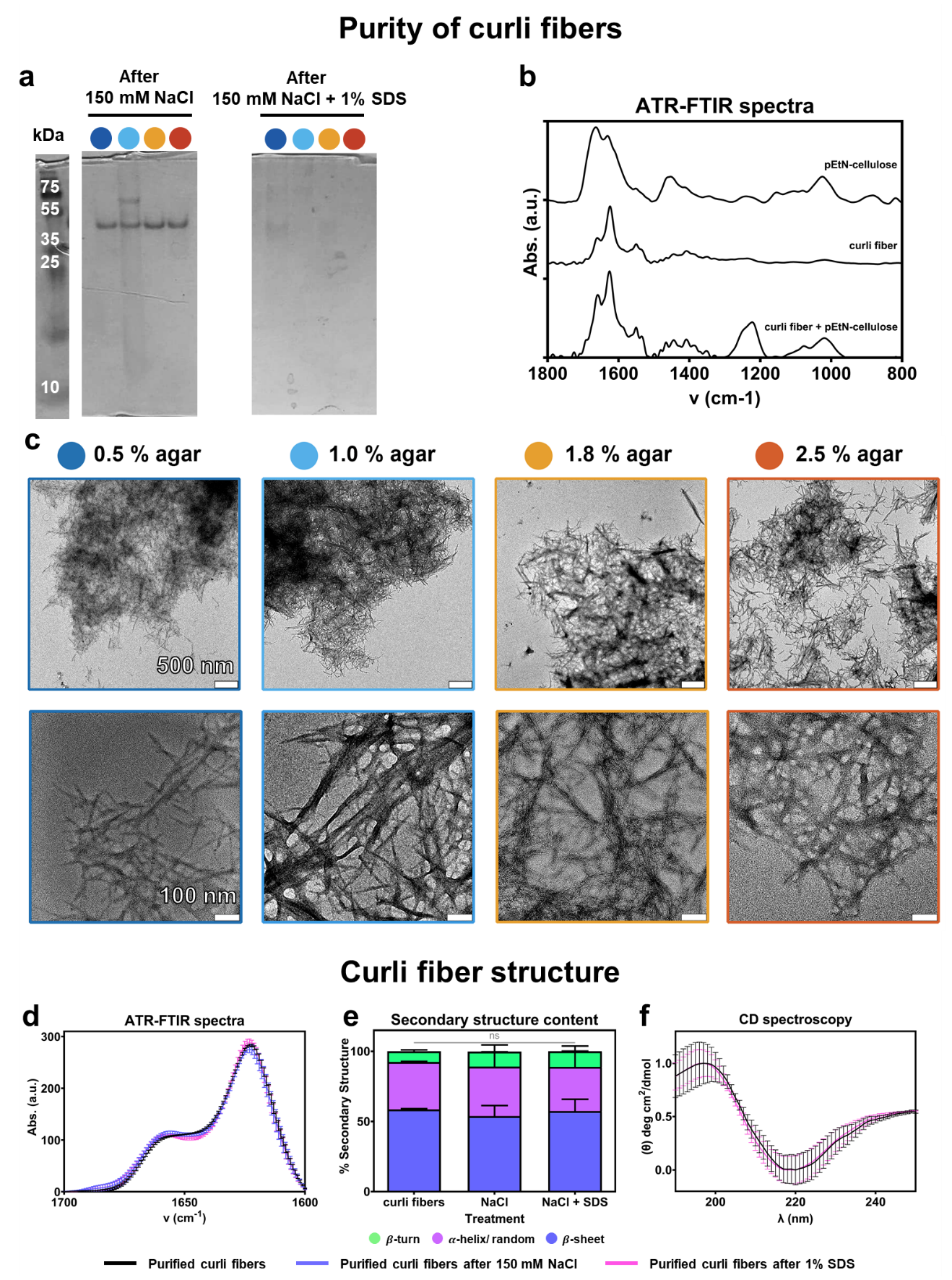

Figure S3 Purity and integrity of extracted curli fibers. (a) 12 % acrylamide SDS-PAGE for each sample. The first four lanes depicts the samples from the purification process for each condition, after the step of ice incubation with 150 mM NaCl 10 mM Tris-HCl buffer pH 7.4 (1). The second four lanes depict the samples of the purification process for each condition, after 1% SDS treatment (2). (b) FTIR spectra of purified *E. coli* cellulose, purified curli fibers and the purified mixture of both *E. coli* cellulose and curli fiber. N= 3 (c) Electron microscopy images of an overview of the purified fiber solution (upper panel, scale bar= 500 nm), and a more detailed image of the fibers purified from biofilms grown on salt-free LB-agar substrates (lower panel, scale bar= 100 nm) containing 0.5 %, 1.0 %, 1.8 % and 2.5 % agar. N= 3- 6 (d) ATR-FTIR spectra of purified curli fibers (black line), the purified curli fibers after incubation in ice with 150 mM NaCl 10 mM Tris-HCl buffer pH 7.4 (blue line) and the purified curli fibers after incubation at 37 °C in 1% SDS (pink line). (e) Secondary component content of the curli fibers. The data was acquired through band fitting of the spectra in panel (d) (for details please see the experimental section). For statistical analysis One-way ANOVA test was used (p<0.05, *| ns = non-significant) with a Dunnett’s post-test for multiple comparisons (alpha=0.05) comparing all samples against the purified curli fibers. There was no significant differences between the conditions tested. (f) CD spectra of purified curli fibers (black line), and the purified curli fibers after incubation at 37 °C in 1% SDS (pink line). N= 3

#### 4. ζ-Potential of purified curli fibers

4.1. **Experimental section**
     4.1.1. ζ-Potential measurements

10 µL of purified fibers were diluted to a final volume of 700 µL of milliQ water or 50 mM glycine buffer pH 8.0, used in the ThioT fluorescence experiments. DTS 1070 polystyrene capillary cuvettes were used for the measurements. Samples were equilibrated for 20 seconds at 25 °C. The experiment was carried out in a Malvern Zetasizer (Zetasizer Nano ZS, Malvern, Worcestershire, UK). Measurements of each sample was done by triplicate.

4.2. **Results**

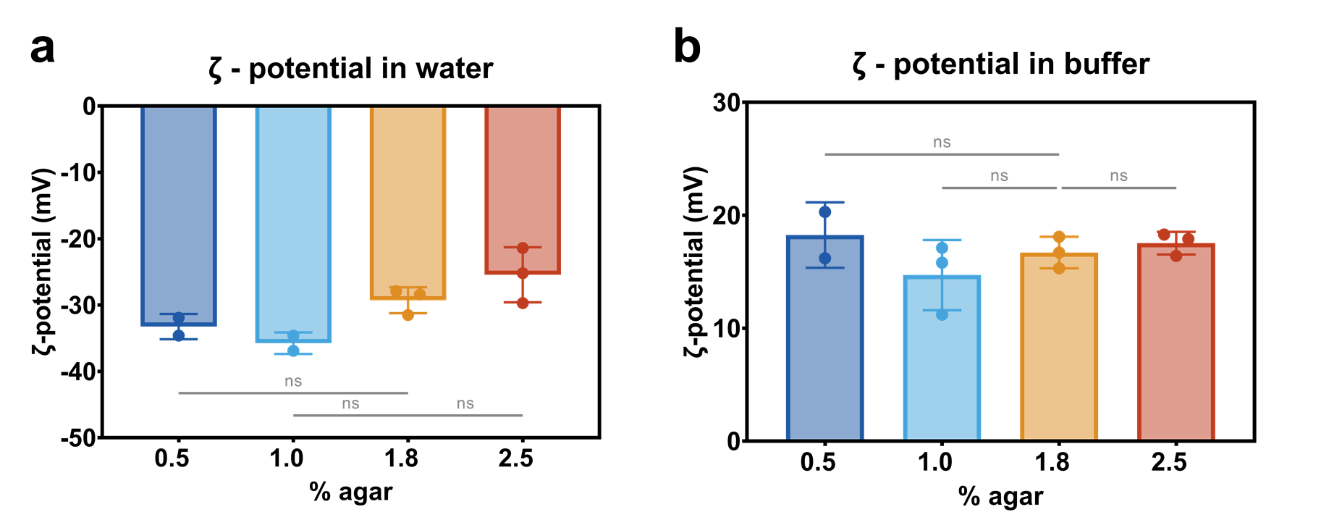

Figure S 4 ζ-Potential value of purified curli fibers from *E. coli* W3110 biofilms in (a) milliQ water and (b)50 mM glycine buffer pH 8.0. For statistical analysis On-way ANOVA was used (p<0.1, * | ns = non-significant) with a Dunnett’s post-test for multiple comparisons, comparing all conditions to the 1.8 % agar substrate.

#### 5. FT-IR spectra of the purified curli fibers

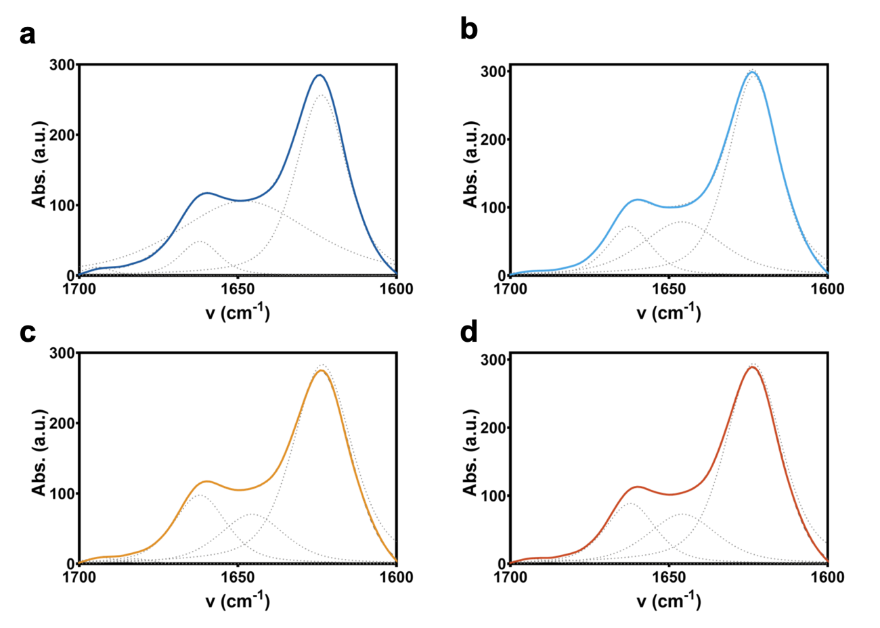

Figure S5 Amide I’ spectra of curli fibers purified from *E. coli* biofilms. Area-normalized FTIR absorbance spectra in the Amide I’ region of the different samples. The spectra were curve fitted (dashed lines) using the number of spectral components identified by second derivative on the Fourier self-deconvoluted spectra. N= 4.

#### 6. Spectral phasor plots

In the spectral phasor transformation each spectrum will represent a single point in the phasor plot. The total wavelength range in which the spectrum was acquired will be plotted starting from 0 degrees until 360 degrees. The spectral shift is related to the angular position in the phasor plot, while the width of the spectrum is related to the distance from the center as exemplified in Figure S3.

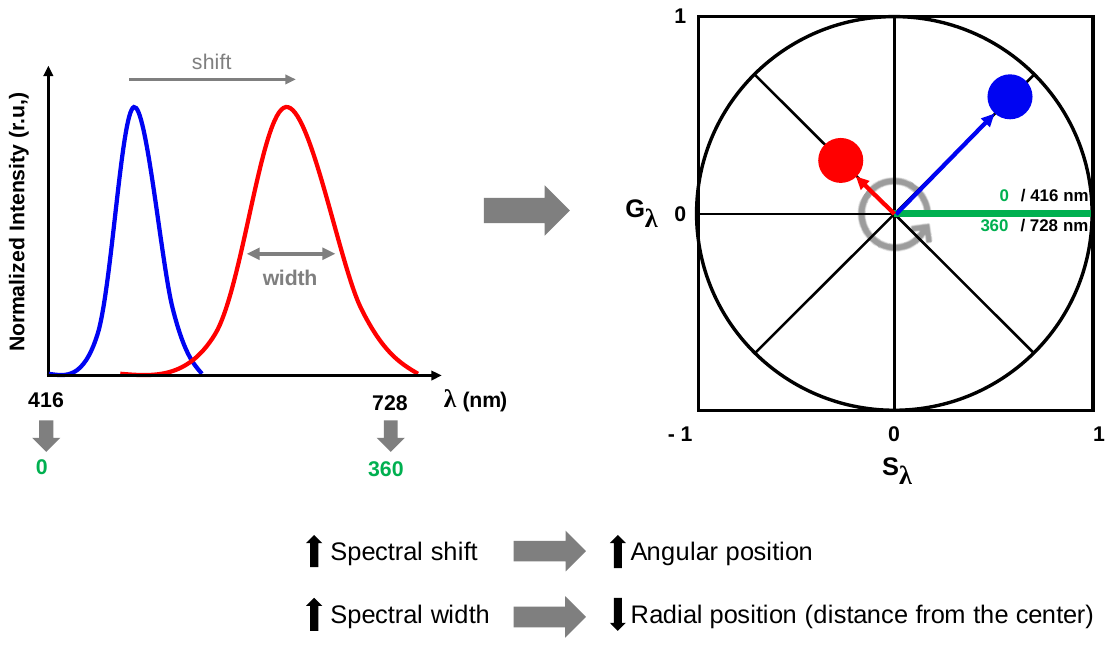

Figure S6 Spectral phasor transformation. In the phasor plot, the wavelength increase is visualized counter-clockwise (see grey arrow at the center) beginning and ending in the green line at (1,0). For that reason, the point corresponding to the red spectrum (higher wavelength) appears at a higher angular position from the origin compared to the blue point. The radial position of the points in the phasor plot is related with the spectral width; with the broader spectra appearing towards the center of the plot (0, 0).

#### 7. FLIM phasor plots

In FLIM phasor plots, all the single exponential lifetimes will lie on the “universal circle”, defined as the semicircle going from (0,0) to (1,0), with radius 0.5. Point (1,0) corresponds to $\tau=0$, and point (0, 0) to $\tau=\infty$. Then, longer lifetimes appear towards the origin (x=0) and shorter lifetimes towards the bottom right intersection with the x axis (x=1). Complex decays will fall inside the universal circle. In this manner, the phasor plot allows a quick interpretation of the data by single inspection, without the need to use models to fit the data, as exemplified below in Fig. S3.

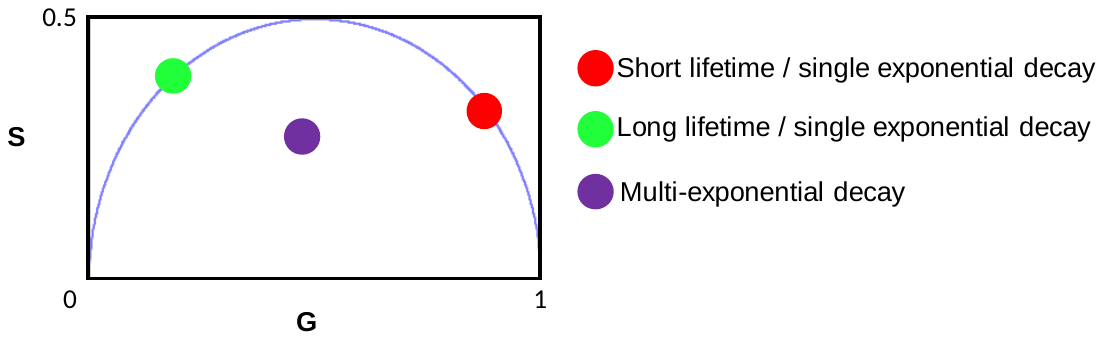

Figure S7 Lifetime phasor plot. Single-exponential lifetimes will fall in the universal circle, decreasing in lifetime going from left to right (green and red dots). Complex decays will fall inside the universal circle. In this manner, lifetimes changes and the nature of the lifetime decay (single or multi-exponential) can be easily observed by single inspection of the plot, and without the need of fitting the data.

#### 8. Nile red FLIM analysis

Quantification of the change in lifetime of nile red in the different fibers was achieved using the two-cursor analysis of the SimFCS software (Ranjit et al 2018).^3^ We positioned the red and green cursors as indicated in Fig. S4a, to calculate the histogram for the pixel distribution along the line. In this way, we can quantify how the pixel clouds move towards longer or shorter lifetimes for each sample. The average value for each histogram (± SD) is plotted in Fig S4b. A better way to observe the differences in the histograms is by calculating the center of mass of the distributions^4^ as showed in Figure S4c. Center of mass is calculated as:

$$CM=\frac{\Sigma_{i=0}^{i=1}F_{i}i}{\Sigma_{i=0}^{i=1}F_{i}}$$

where $F_{i}$ is the number of pixels at a given fraction ($i$).

It can be clearly observed that there are significative differences between the samples.

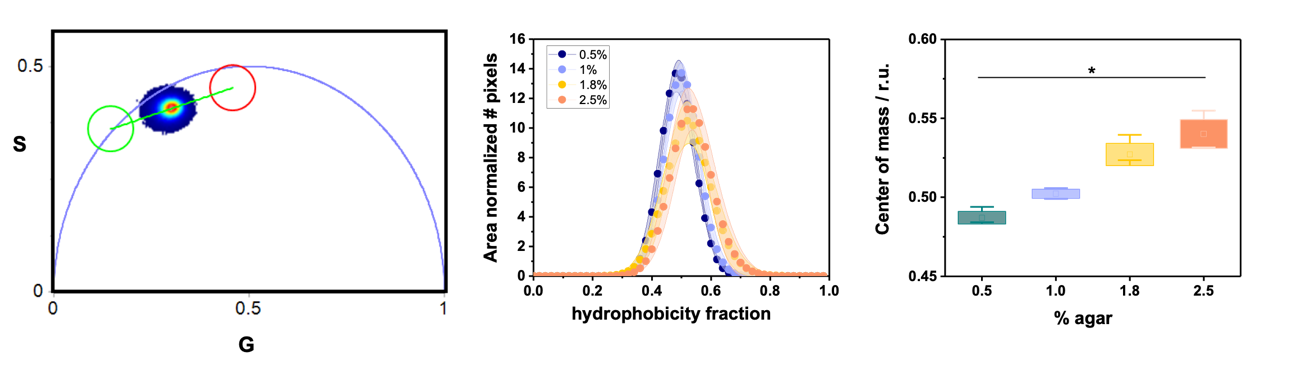

Figure S8 Statistical quantification of the data shown in Figure 7. (a) Phasor plot and two cursors linear analysis. (b) Histograms of the hydrophobicity fraction of the purified fibers showed as mean values ± standard deviation. (c) Center of mass of the hydrophobicity fraction of each fiber. Squares indicate the mean values and boxes sizes the standard deviation. The wiskers indicate the maximum and minimum values of the distributions. n=5 per condition, differences are significative: p<0.05, * ANOVA and Tukey post-test analysis.

#### 9. Raw spectra of the quenching experiments and Ksv values

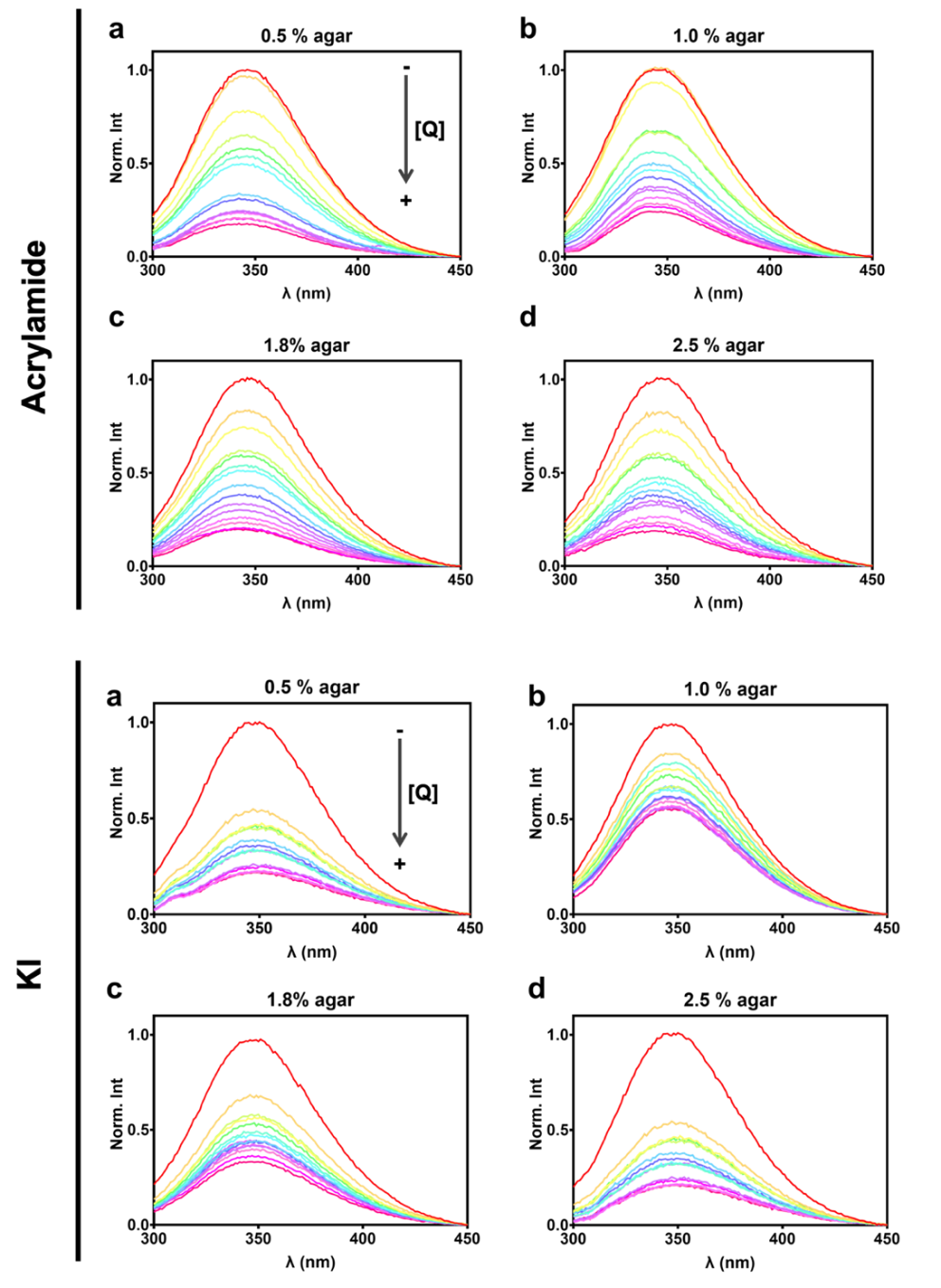

Figure S9 Effect of the quencher (acrylamide or KI) in the spectra emission of the fibers. Intensity curves at increasing quencher concentration (0.0 – 0.4 M).

Table S2 Stern-Volmer constants (M^− 1^) for the different purified curli amyloid fibrils and NATA

|  | **0.5 % Agar** | **1.0 % Agar** | **1.8 % Agar** | **2.5 % Agar** | **NATA** |
| --- | --- | --- | --- | --- | --- |
| **Acrylamide** | 12.66 $\pm$ 0.66 | 10.29 $\pm$ 0.61 | 11.23 $\pm$ 0.41 | 12.60 $\pm$ 0.41 | 60.88 $\pm$ 3.70 |
| **KI** | 5.36 $\pm$ 0.43 | 4.28 $\pm$ 0.20 | 5.84 $\pm$ 0.35 | 5.64 $\pm$ 0.61 | 10.67 $\pm$ 0.40 |

#### 10. Complete fiber thermal denaturation ramp (ATR-FTIR analysis)

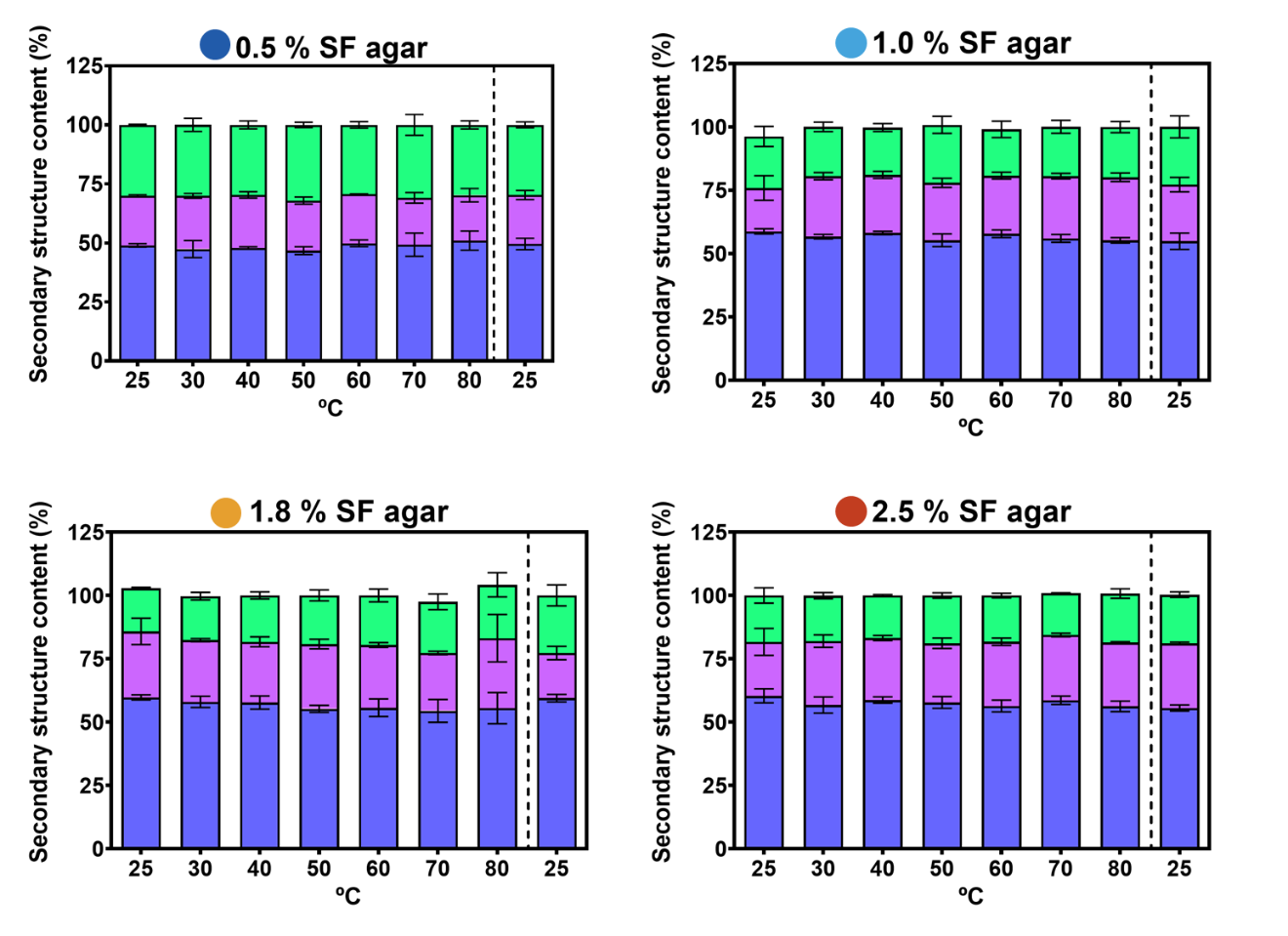

Figure S10 Thermal stability of fibers purified from biofilms grown on substrates of different water contents. Secondary structure contents by analysis of FTIR spectra of curli fibers treated with the complete temperature ramp (25 – 80 °C). The data after the dashed line represent the secondary structure contents of fibers after a sudden decrease of temperature (from 80 – 25 °C). N=3 fiber solutions per condition.

#### 11. Micro-indentation of E. coli W3110 biofilms

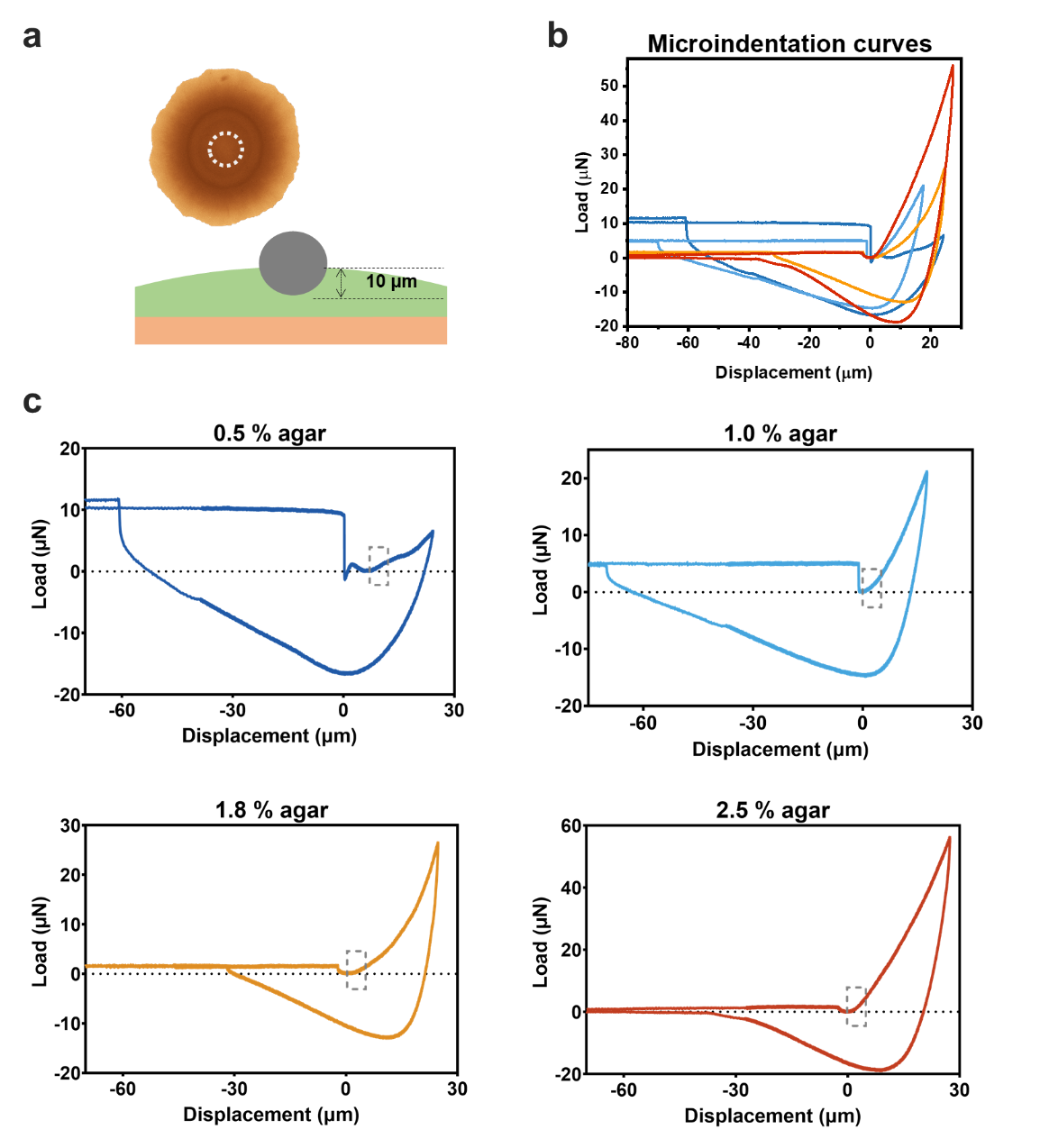

Figure S11 Micro-indentation of the *E. coli* W3110 biofillms. (a) Scheme of how the experiment was carried out. The bead used in the tip has a diameter of 50 µm. (b) Representation of full micro-indentation curves for each condition (each curve is representative of the data batch). (c) We represented each micro-indentation curve separately with the region used for fitting encased in the grey square.

#### 12. Fiber structure/ fiber function relationship

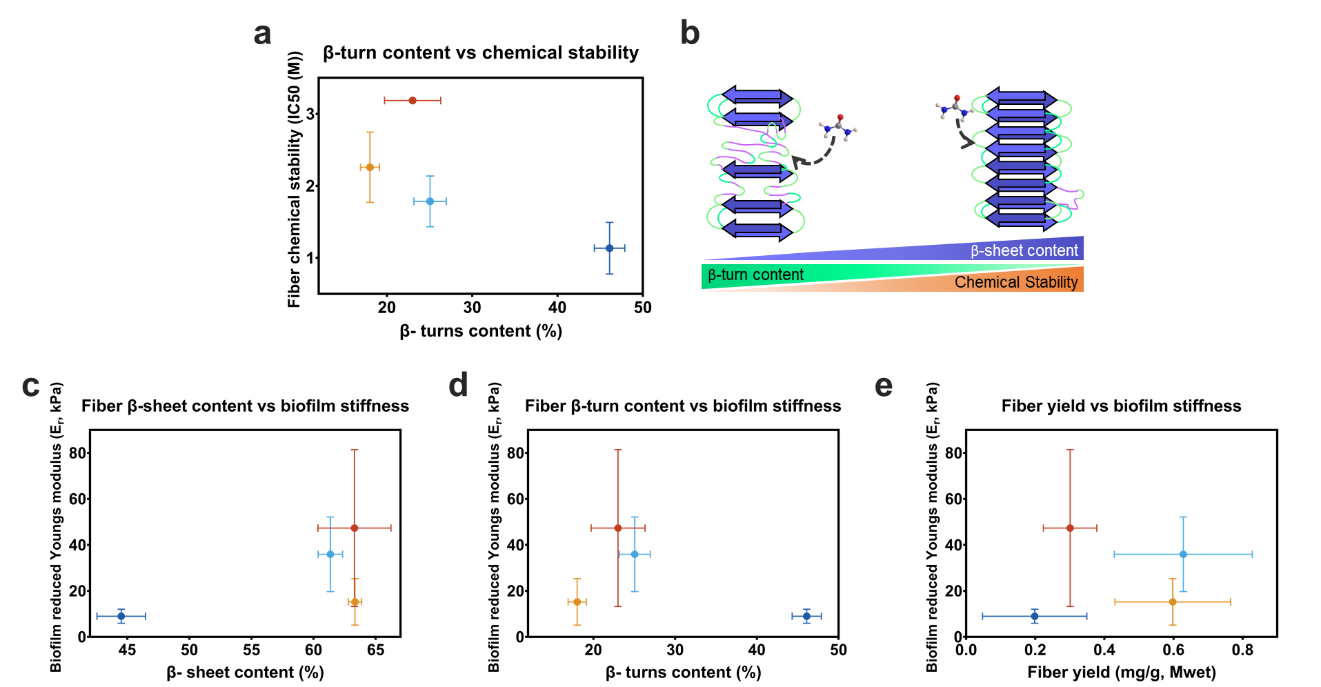

Figure S12 Fiber structure/ fiber function relationship. (a) Relationship between the fibers’ β-turn content and their chemical stability. (b) Scheme depicting how the secondary structure of the fiber influence in the chemical stability of the fiber. (c-d) Relationship between the fibers’ secondary structure composition and the biofilm stiffness. (e) Relationship between the fibers’ yield and the biofilm stiffness. Color code: 0.5 % agar content (blue), 1.0 % agar content (light-blue), 1.8 % agar content (orange), 2.5 % agar content (red).

**References**

1.	Ziege, R. *et al.* Adaptation of Escherichia coli Biofilm Growth, Morphology, and Mechanical Properties to Substrate Water Content. *ACS Biomater. Sci. Eng.* **7**, 5315–5325 (2021).

2.	Bertasa, M. *et al.* A study of non-bounded/bounded water and water mobility in different agar gels. *Microchem. J.* **139**, 306–314 (2018).

3.	Ranjit, S., Malacrida, L., Jameson, D. M. & Gratton, E. Fit-free analysis of fluorescence lifetime imaging data using the phasor approach. *Nat. Protoc.* **13**, 1979–2004 (2018).

4.	Malacrida, L. & Gratton, E. LAURDAN fluorescence and phasor plots reveal the effects of a H2O2 bolus in NIH-3T3 fibroblast membranes dynamics and hydration. *Free Radic. Biol. Med.* **128**, 144–156 (2018).
